# A vagal gut-brain axis for feeding induced gastric secretion

**DOI:** 10.64898/2026.03.01.708884

**Authors:** Zhuang Wang, Qihong Tang, Kai Li, Jing Li, Junhui Mou, Xiaohan Sun, Yujing Wei, Yuanyuan Chen, Peng Guo, Lei Jin, Benlong Liu, Bailong Xiao, Shumin Duan, Wei Dai, Jinfei D Ni

## Abstract

Feeding begins with food-seeking and ends with nutrient absorption and waste elimination. Although meal-related sensory cues are known to shape feeding behavior, how they engage neural circuits to control digestion remains unclear. We dissected vagal gut-brain-gut pathways that mediate meal-induced gastric secretion. Vagal afferent neurons innervating the stomach were necessary and sufficient for secretion. Among those, *Glp1r* and *Sst* neurons robustly drove acid, pepsin, and gastrin secretion, whereas *Oxtr* and *Vip* neurons had little effect. *Piezo2* was expressed in *Glp1r* mechanosensory afferents, and conditional deletion of Piezo channels attenuated distension-evoked secretion. On the efferent limb, genetically defined vagal motor neuron subsets differentially recruited secretory outputs: *Cck* neurons broadly increased acid, pepsin, and gastrin, whereas *Grp* neurons preferentially drove acid secretion. Finally, enteric circuit dissection revealed modular output control: *Calb2* enteric neurons potently drove secretion with minimal motility effects, *Grp* enteric neurons coupled secretion to motility, and *Cysltr2* enteric neurons selectively modulated acid output. Together, these results define a cell-type resolved vagal-enteric architecture for gastric control and show that secretory and motor programs can be segregated across dedicated enteric neuron subtypes.

## INTRODUCTION

Feeding begins with food-seeking and ends with nutrient absorption and energy replenishment. Food-related sensory cues regulate each step of feeding by engaging a complex, hierarchical neuronal network. The sight, smell, and taste of food shape food-seeking behavior and food preference. Once ingested, the physical presence and composition of a meal modulate gastrointestinal physiology to promote digestion and absorption. Signals arising from ingested and partially digested food also provide anticipatory cues for energy homeostasis and contribute to satiety^1–7^. Among these coordinated responses, gastric secretion—acid release and pepsin activation—is an early effector that must be precisely timed with food entry into the stomach^8^.

Vagal sensory neurons, whose cell bodies reside in the nodose ganglia, monitor internal signals from multiple organs, including the gastrointestinal tract^9^. These neurons are heterogeneous in terminal morphology, sensory properties, and gene expression^10–12^. Within the gut, vagal afferents form specialized endings that detect luminal chemicals and mechanical stimuli generated by a meal^13^. Activated vagal afferents relay food-related information to the brainstem and engage vago-vagal reflexes that modulate postprandial physiology and behavior—gastrointestinal secretion, motility, metabolic responses, and satiety among them^9,14–20^.

Vagal motor neurons in the DMV form the efferent arm of gut–brain–gut reflexes and provide the dominant parasympathetic outflow to the stomach. These preganglionic neurons project via the vagus nerve to the stomach and small intestine to control secretion, motility and gut immunity^18,21,22^. Meal-evoked vagal afferent signals are integrated in the NTS and rapidly routed to the DMV through excitatory and inhibitory synapses, enabling sensory cues and distension to shape gastric output. Importantly, the DMV is not a uniform pool of “generic” parasympathetic motor neurons: recent single-nucleus transcriptomic and genetic-access studies resolved multiple transcriptionally distinct *Chat* DMV subtypes, including populations marked by neuropeptides such as Cck and Pdyn that show highly selective stomach targeting and differential recruitment of enteric effector modules^23^.

While vago–vagal reflexes are well established physiologically, how defined sensory modalities and organ segments signal feeding to drive gastric secretion, how afferent signals are parsed into specific vagal motor outputs, and which enteric neurons execute the secretomotor response remain incompletely understood. Here we address these gaps by linking mechanical and nutrient cues generated by a meal to identified vagal afferents, mapping the efferent DMV subsets that promote secretion, and pinpointing an enteric relay that link brainstem commands to the gastric epithelium. This work defines a vagal gut-brain-enteric axis for feeding-induced gastric secretion and clarifies how early post-ingestive signals are translated into coordinated steps of food digestion.

## RESULTS

### Food induced gastric secretion

Food ingestion initiates a digestive program that is tightly orchestrated by the nervous system^24,25^. Among these coordinated processes, gastric secretion is a critical step that initiates food digestion by supplying gastric acid and digestive enzymes^15,26^. We therefore asked whether ingested food could promote gastric secretion. We gavaged mice with liquid food and measured intragastric pH and pepsin activity at three time points over the subsequent 30 minutes **(Figure S1A)**. Liquid food significantly promoted gastric secretion, as indicated by a time-dependent reduction in intragastric pH (proton secretion) and an increase in pepsin activity (**Figure S1B**).

Two types of sensory information are generated by ingested food: mechanical distension and nutrient-related chemical stimuli. We next asked which type of sensory information induces gastric secretion. To test the effect of mechanical distension, a latex balloon was inserted into different segments of the GI tract (esophagus, stomach, duodenum, and distal colon) to deliver distension. When balloon distension was administered in the stomach, we observed a dose-dependent reduction in gastric pH and an increase of pepsin activity (**Figure S1C and Figure 1A-B**). By contrast, mechanical distention in other segments of the gastrointestinal tract did not lead to a significant increase in gastric secretion. (**Figure 1C and Figure S1D-E**). These results indicate that food-generated mechanical distension in the stomach powerfully drives gastric secretion.

**Figure 1.**
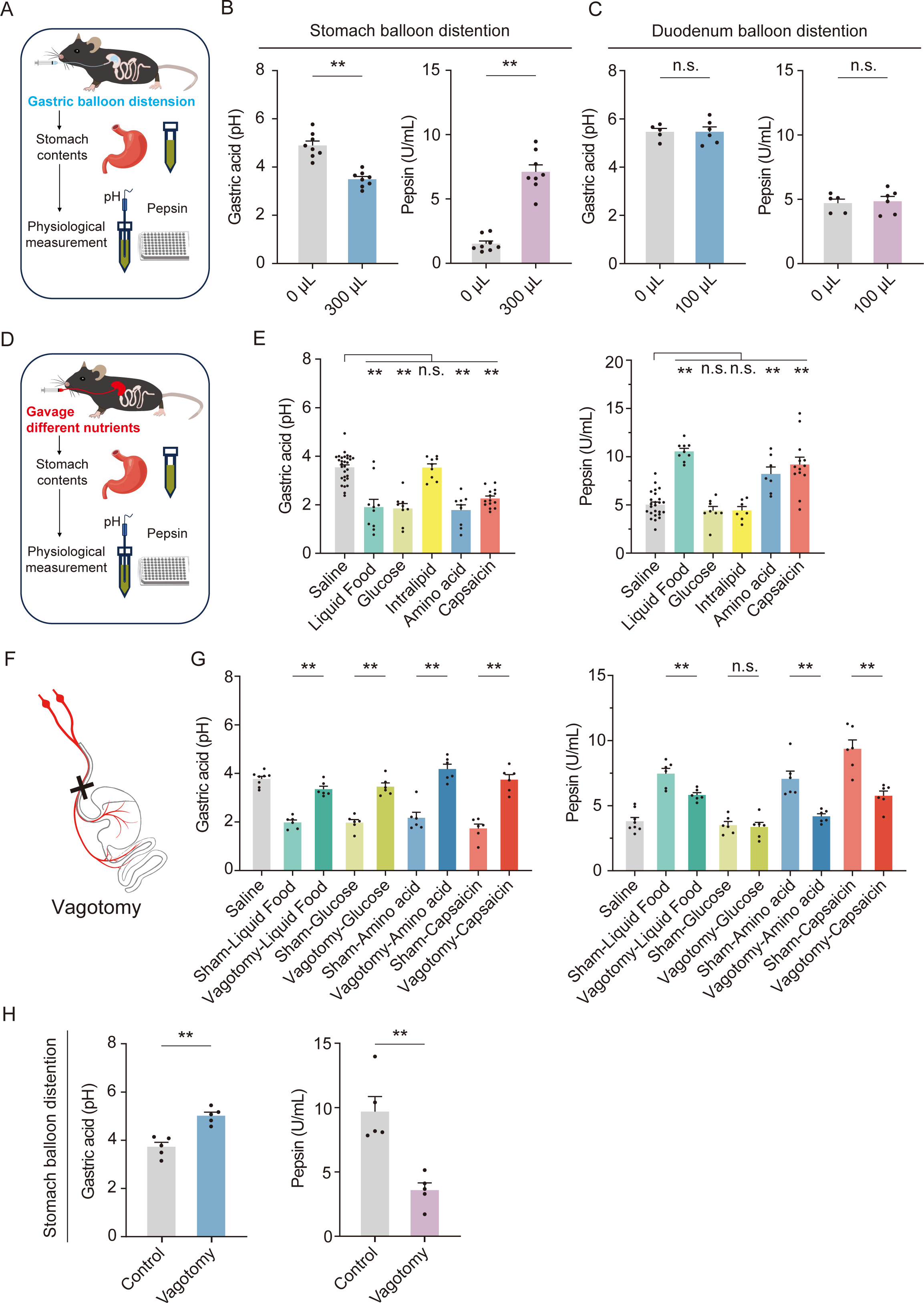
Gastric secretion induced by mechanical and nutrient-related chemical stimulation**. (A)** Schematic of gastric mechanical distension and gastric secretion measurements. **(B-C)** Gastric acid (left) and pepsin (right) secretion evoked by mechanical distension of the stomach (B) or duodenum (C). Data are mean ± SEM. Unpaired two-tailed t test. n.s., not significant; **P < 0.01. Each dot represents data from one mouse. **(D)** Schematic of oral gavage–based nutrient stimulation and gastric secretion measurements. **(E)** Effects of nutrient stimuli on gastric acid (left) and pepsin (right) secretion. Data are mean ± SEM. One-way ANOVA with Tukey’s multiple-comparisons test. n.s., not significant; **P < 0.01. Each dot represents data from one mouse. **(F)** Schematic of subdiaphragmatic vagotomy. **(G)** Subdiaphragmatic vagotomy reduces nutrient-evoked gastric acid (left) and pepsin (right) secretion. Data are mean ± SEM. One-way ANOVA with Tukey’s multiple-comparisons test. n.s., not significant; **P < 0.01. Each dot represents data from one mouse. **(H)** Subdiaphragmatic vagotomy reduces distension-evoked gastric acid (left) and pepsin (right) secretion. Data are mean ± SEM. Unpaired two-tailed t test. **P < 0.01. Each dot represents one mouse; n = 5 mice per group.

To test the effect of chemical signals on gastric secretion, we gavaged mice with different nutrient-related chemicals as well as saline (negative control) or liquid food (positive control) (**Figure 1D**). Nutrient-related chemicals differentially affected gastric secretion: whereas an amino acid mixture increased the secretion of gastric acid and pepsin, glucose increased gastric acid secretion without measurably altering pepsin secretion. In contrast, intralipid did not affect either gastric acid or pepsin secretion (**Figure 1E**). Notably, the TRPV1 agonist capsaicin^27^ also potently stimulated gastric secretion, indicating a potential sensory-autonomic reflex involving TRPV1^+^ sensory neurons (**Figure 1E**). Infusing liquid food into the duodenum leads to only a modest increase in gastric secretion (**Figure S1F**), indicating that the stomach is the major chemical sensing site for food induced gastric secretion.

### Vagal afferent pathways for gastric secretion

To test whether the vago-vagal reflex is essential for food-induced gastric secretion, we performed subdiaphragmatic vagotomy to ablate vagal innervation to the stomach and intestine (**Figure 1F**) and then administered nutrient stimuli or mechanical distension. Subdiaphragmatic vagotomy significantly blunted both nutrient-and distension-induced gastric acid and pepsin secretion (**Figure 1G-H)**. However, this procedure eliminates both vagal sensory (afferent) and motor (efferent) pathways linking the gut to the brain^28^.

To isolate the contribution of vagal sensory pathways, we chemogenetically activated vagal sensory neurons innervating the stomach or duodenum. Specifically, we injected a retrograde AAV expressing Cre recombinase intramurally into the corresponding segment of the gastrointestinal tract, and in the same animals injected a Cre-dependent hM3Dq^29^ AAV into the nodose ganglion (**Figure 2A-B**). Control mice received the same peripheral AAV-Cre but a Cre-dependent tdTomato in the nodose ganglion. Compared with controls, activating stomach-innervating vagal sensory neurons led to robust increases in gastric acid and pepsin secretion (**Figure 2C-D**), whereas activating duodenum-innervating vagal sensory neurons produced little increase in gastric secretion (**Figure 2C-D**). We also measured the impact of vagal sensory neuron activation on gastrin, a peptide hormone produced by antral G cells that stimulates gastric acid secretion and promotes mucosal growth^30–34^. Consistent with the effects on acid and pepsin, activating stomach-innervating—but not duodenum-innervating—vagal sensory neurons significantly increased gastrin release (**Figure 2E).**

**Figure 2.**
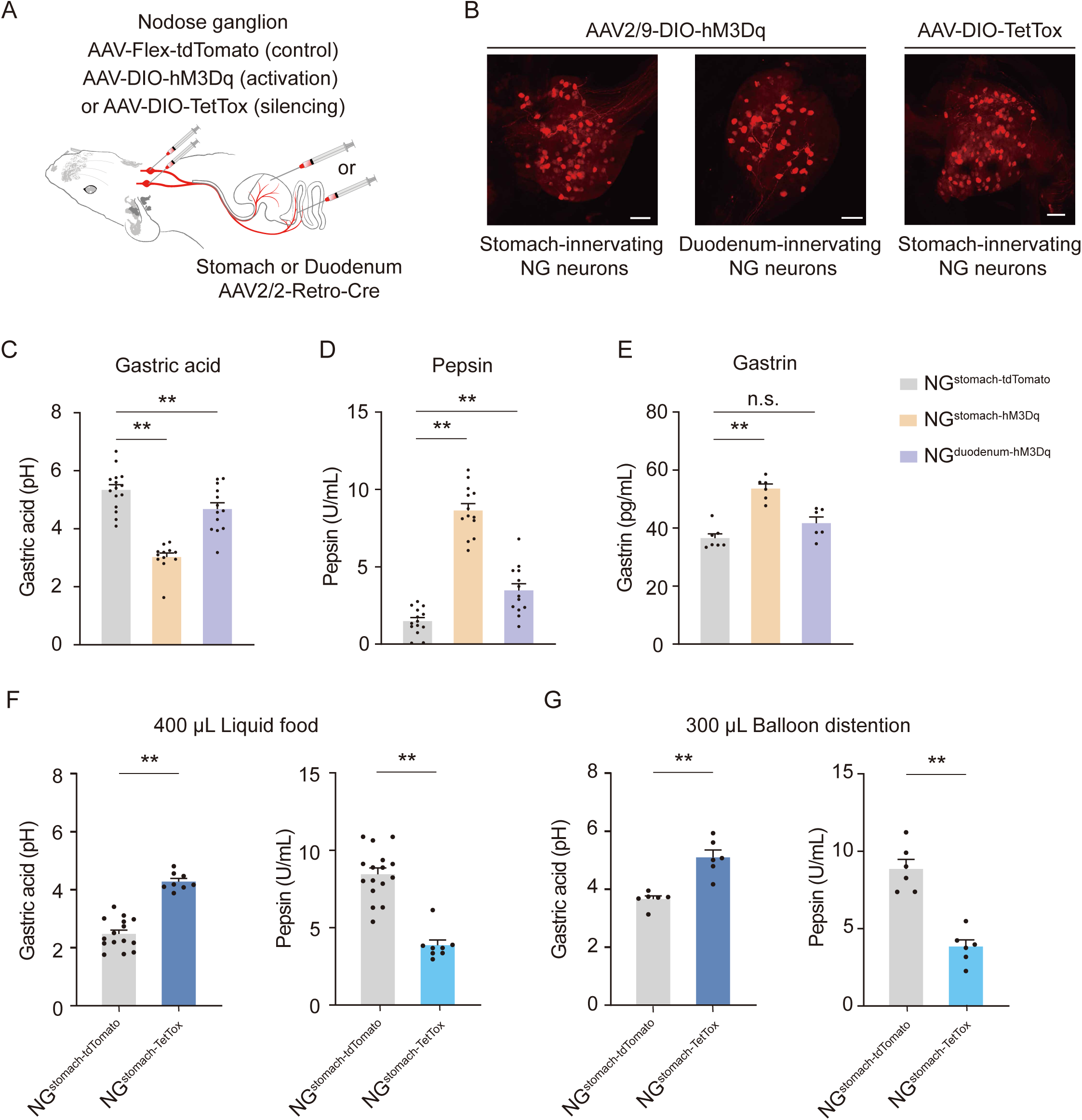
Vagal sensory neurons mediate gastric secretion. **(A)** Schematic of AAV-mediated gene delivery to vagal sensory neurons that project to the stomach or duodenum. **(B)** Whole-mount immunostaining of the nodose ganglion showing AAV-labeled vagal sensory neurons expressing the excitatory DREADD hM3Dq or the synaptic silencer TetTox. Scale bar, 100 μm. **(C–E)** Chemogenetic activation of stomach-innervating vagal sensory neurons increases gastric acid (c), pepsin (d), and gastrin (e) secretion. In contrast, activation of duodenum-innervating vagal sensory neurons has little effect on gastric secretion. Data are mean ± SEM. One-way ANOVA with Tukey’s multiple-comparisons test. n.s., not significant; **P < 0.01. Each dot represents data from one mouse. **(F–G)** Silencing stomach-innervating vagal sensory neurons reduces gastric acid and pepsin secretion evoked by liquid food (F) or mechanical distension (G). Data are mean ± SEM. Statistical significance was determined using Unpaired two-tailed t test. **P < 0.01. Each dot represents data from one mouse.

To test the requirement of vagal sensory neurons for feeding-induced gastric secretion, we selectively silenced stomach-innervating vagal sensory neurons by expressing tetanus toxin light chain (TetTox) ^35,36^ using the same viral genetic strategy for vagal sensory neuron activation (**Figure 2B, right panel**). Mice were administrated with a liquid meal by oral gavage and intragastric pH and pepsin activity were measured 30 minutes later. Compared to control animals (expressing tdTomato in vagal sensory neurons), silencing stomach-innervating vagal sensory neurons resulted in higher intragastric pH (reduced proton secretion) and lower pepsin activity (**Figure 2F**). Silencing stomach-innervating vagal sensory neurons also dampened gastric secretion induced by mechanical distension (**Figure 2G**). Together, vagal sensory neuron activation and silencing experiments indicate that stomach-innervating vagal sensory neurons play a major role in food-induced gastric secretion.

Vagal sensory neurons innervating the gastrointestinal tract comprise physiologically and molecularly distinct subpopulations that can be accessed with different Cre driver lines^15^. We used the following four Cre lines marking subsets of gut-innervating vagal sensory neurons: *Glp1r-Cre*, *Oxtr-Cre*, *Vip-Cre*, and *Sst-Cre* (**Figure 3A**)^13,15^. To test which subsets are sufficient to drive gastric secretion, we injected AAV expressing Cre-dependent hM3Dq into the nodose ganglion of each Cre line. AAV expressing Cre-dependent tdTomato was injected as control (**Figure 3B**). Activating *Glp1r-Cre*^+^ vagal sensory neurons, which include mechanosensory neurons innervating both stomach and duodenum^13,15^, produced significant increases in gastric acid and pepsin secretion (**Figure 3C-D, Figure S3A-C)**. Activating *Oxtr-Cre*^+^ vagal sensory neurons, which are mechanosensitive and predominantly innervate the duodenum while only sparsely innervate the stomach, did not induce gastric secretion (**Figure 3E-F, Figure S3D-F**), although both *Glp1r-Cre*^+^ and *Oxtr-Cre*^+^ mechanosensitive neurons negatively regulate feed (**Figure S3G-H**). *Vip-Cre*^+^ neurons, which innervate the villi of small intestine^15^ and the gastric corpus (**Figure 3G, Figure S4A-C)** did not evoke gastric secretion **(Figure 3H)** nor alter feeding^15^ **(Figure S4G)**. In contrast, activating *Sst-Cre*^+^ vagal sensory neurons increased gastric acid and pepsin secretion, although the effect on gastrin release was smaller (**Figure 3J**). These neurons also had a modest positive effect on feeding behavior (**Figure S4H**). We analyzed the projection pattern of *Sst-Cre*^+^ vagal sensory neurons in the gastrointestinal tract and found that they specifically innervate the villi of the stomach antrum—the same region enriched with gastric chemosensory enteroendocrine cells^37^ (**Figure 3I, and Figure S4D-F**).

**Figure 3.**
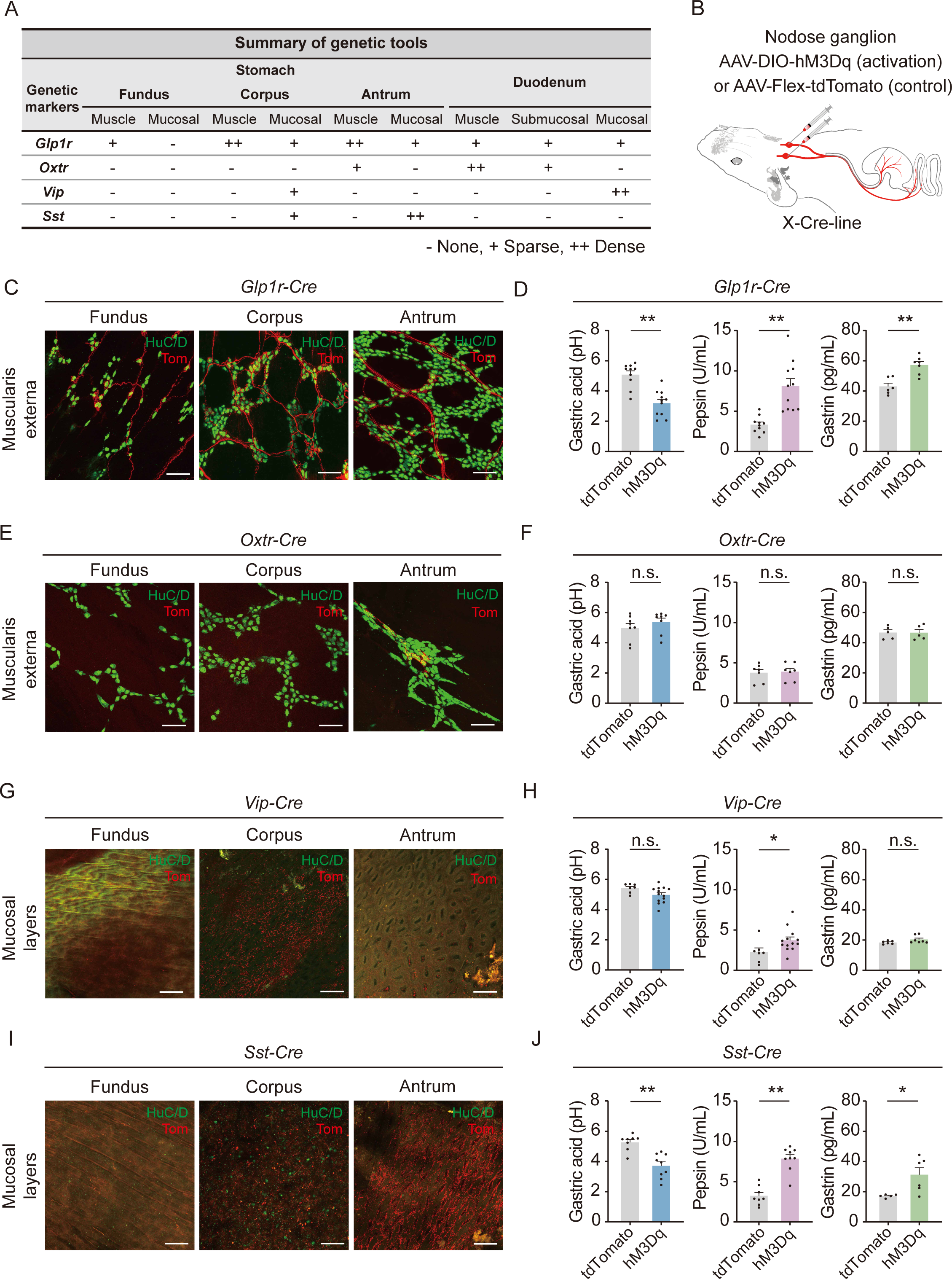
Distinct subtypes of stomach-innervating vagal sensory neurons differentially regulate gastric secretion. **(A)** Summary of gastric versus duodenal projection patterns across four Cre-driver mouse lines. **(B)** Schematic of nodose ganglion injections. **(C)** *Glp1r* vagal sensory neurons densely innervate the gastric muscularis externa. Scale bar, 100 μm. **(D)** Chemogenetic activation of *Glp1r* vagal sensory neurons increases gastric acid (left), pepsin (middle), and gastrin (right) secretion. Data are mean ± SEM. Unpaired two-tailed t test. **P < 0.01. Each dot represents data from one mouse. **(E)** *Oxtr* vagal sensory neurons sparsely innervate the gastric muscularis externa. Scale bar, 100 μm. **(F)** Chemogenetic activation of *Oxtr* vagal sensory neurons does not impact gastric acid (left), pepsin (middle), or gastrin (right) secretion. Data are mean ± SEM. Unpaired two-tailed t test. n.s., not significant. Each dot represents data from one mouse. **(G)** Innervation pattern of *Vip* vagal sensory neurons in gastric mucosal layers. Scale bar, 100μm. **(H)** Chemogenetic activation of *Vip* vagal sensory neurons does not significantly change gastric acid (left) or gastrin (right) secretion and elicits a modest increase in pepsin secretion (middle). Data are mean ± SEM. Unpaired two-tailed t test. n.s., not significant; *P < 0.05. Each dot represents data from one mouse. **(I)** Innervation pattern of *Sst* vagal sensory neurons in gastric mucosal layers. Scale bar, 100μm. **(J)** Chemogenetic activation of *Sst* nodose ganglion neurons increases gastric acid (left), pepsin (middle), and gastrin (right) secretion. Data are mean ± SEM. Unpaired two-tailed t test. *P < 0.05; **P < 0.01. Each dot represents data from one mouse.

### Piezo channels participate in gastric secretion

Given that both mechanical distension and mechanosensitive vagal sensory neurons robustly promote gastric secretion (**Figure 1B and Figure 3C-D**), we sought to identify mechanosensitive proteins mediating stomach distension-induced secretion. A useful candidate should be enriched in *Glp1r*^+^ vagal sensory neurons^13^ but largely absent from *Vip*^+^ or *Sst*^+^ chemosensory neurons^15^.

We reanalyzed a previously published single-cell sequencing dataset for vagal sensory neurons^16^, focusing on stomach-projecting populations, and performed clustering analysis. This resulted in 16 subsets of stomach-innervating vagal sensory neurons, including subpopulations expressing *Glp1r*, *Oxtr*, *Vip*, or *Sst* (**Figure 4A**). Next, we performed differential gene expression analysis, comparing *Glp1r*-expressing neurons against *Vip*-expressing or *Sst*-expressing populations. Among genes significantly enriched in *Glp1r*^+^ neurons, *Piezo2*, encoding a bona fide mechanosensitive cation channel implicated in multiple sensory processes mediated by vagal sensory neurons^38–40^, emerged as a leading candidate (**Figure 4B and Figure S5A**). Consistent with this expression pattern, *Piezo2-Cre* labeled vagal sensory neurons bearing intraganglionic laminar endings (IGLEs) enriched in the stomach wall but largely spared the small intestine (**Figure 4D-F**).

**Figure 4.**
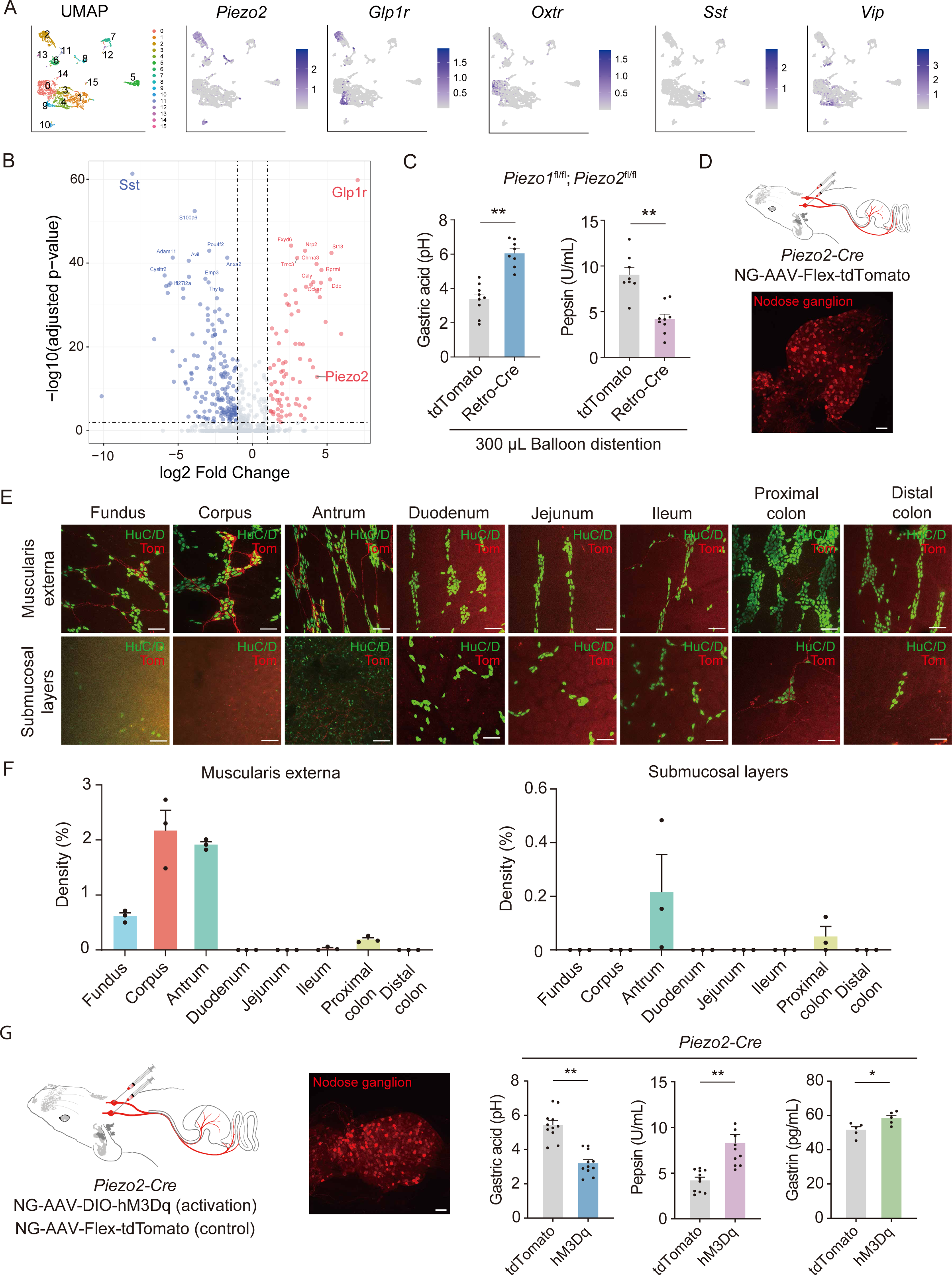
*Piezo2* vagal sensory neurons contribute to gastric secretion. **(A)** UMAP visualization of single-cell transcriptomic data of stomach-innervating vagal sensory neuron, identifying 16 neuronal clusters (left) and feature plots showing expression of *Piezo2*, *Glp1r*, *Oxtr*, *Sst*, and *Vip* (right). **(B)** Volcano plot showing differentially expressed genes between *Glp1r* and *Sst* vagal sensory neuron populations. **(C)** Conditional deletion of *Piezo1* and *Piezo2* in stomach-associated neurons reduces mechanical stimulation–evoked gastric acid (left) and pepsin (right) secretion. Data are mean ± SEM. Unpaired two-tailed t test. **P < 0.01. Each dot represents data from one mouse. **(D)** Schematic of AAV injection into the nodose ganglion of *Piezo2-Cre* mice (top) and representative labeling of *Piezo2* neurons in the nodose ganglion (bottom). Scale bar, 100 μm. **(E–F)** Whole-mount immunostaining showing *Piezo2* vagal sensory neuron terminals across gastrointestinal segments (E) and quantification of terminal distributions (F). HuC/D labels enteric neurons. Scale bar, 100 μm. **(G)** Chemogenetic activation of *Piezo2* vagal sensory neurons increase gastric acid (left), pepsin (middle), and gastrin (right) secretion. Data are mean ± SEM. Unpaired two-tailed t test. *P < 0.05; **P < 0.01. Each dot represents data from one mouse.

To test whether Piezo channels are required for distension-induced gastric secretion, we injected retrograde AAV-hSyn-Cre into the stomach wall of *Piezo1^flox/flox^*; *Piezo2^flox/flox^*mice^41,42^. This resulted in Cre recombinase expression in both stomach-innervating vagal sensory neurons and gastric enteric neurons, but not vagal motor neurons (**Figure S5B**), thereby deleting both Piezo channels in stomach associated sensory neurons. We then subjected these mice and control mice (injected with AAV-tdTomato) to intragastric balloon distension (300 µL, 30 min) and measured intragastric pH and pepsin activity. Compared with controls, deleting Piezo channels blunted distension-induced gastric secretion (**Figure 4C**). Consistent with sufficiency, chemogenetic activation of *Piezo2-Cre*^+^ vagal sensory neurons increased gastric secretion (**Figure 4G**).

### Vagal efferent pathways for gastric secretion

Vagal motor neurons constitute the principal output arm of the vago–vagal reflex. These neurons reside mainly in the dorsal motor nucleus of the vagus (DMV) and the nucleus ambiguus (NA) and can be subdivided into genetically defined populations with distinct projection targets^23,43–45^. The gastrointestinal tract is innervated predominantly by DMV motor neurons^18^, which have been implicated in multiple facets of gastrointestinal physiology, including motility, secretion, and immune regulation^18,22^. Recent single-nucleus transcriptomic and functional mapping studies identified at least seven transcriptionally distinct *Chat* DMV subtypes and supported a labeled-line organization in which specific molecular populations (e.g., *Cck* and *Pdyn*) show selective stomach targeting and differential recruitment of cholinergic versus nitrergic enteric neurons^23^. To identify the DMV motor neuron populations that drive gastric secretion, we examined four *Cre* driver lines—*Cck-Cre*, *Pdyn-Cre*, *Grp-Cre*, and *Trpv1-Cre* (**Figure 5A**).

**Figure 5.**
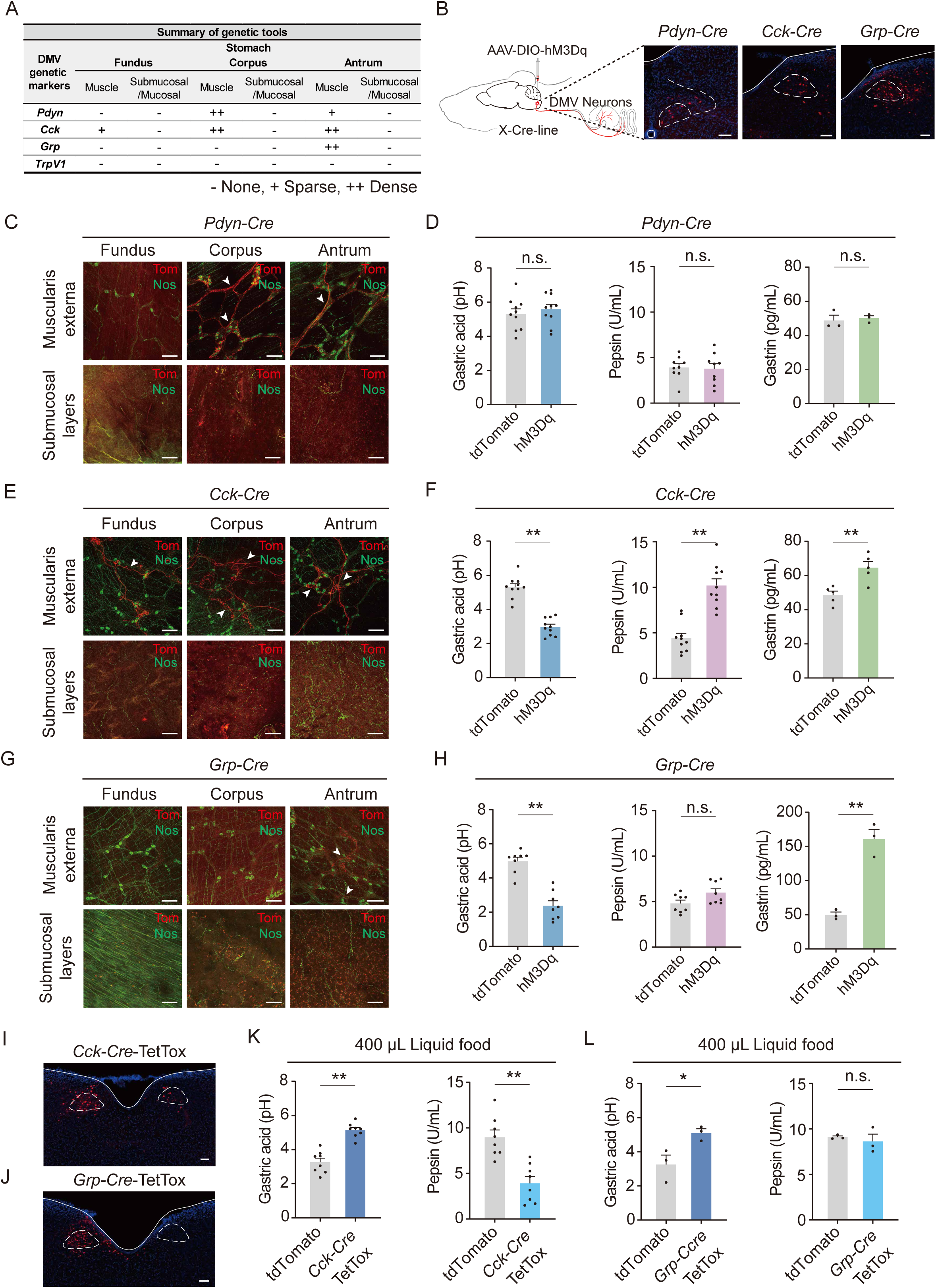
Vagal motor neurons for gastric secretion**. (A)** Summary of gastric projection patterns for four genetically defined dorsal motor nucleus of the vagus (DMV) motor neuron subtypes. **(B)** Schematic of Cre-dependent AAV delivery to the DMV (left) and representative labeling of DMV neurons in *Pdyn-Cre*, *Cck-Cre*, and *Grp-Cre* mice (right). Scale bar, 100 μm. **(C)** Innervation pattern of *Pdyn* DMV neurons in the gastric muscularis externa and submucosal layers. Scale bar, 100 μm. **(D)** Chemogenetic activation of *Pdyn* DMV neurons does not significantly alter gastric acid (left), pepsin (middle), or gastrin (right) secretion. Data are mean ± SEM. Unpaired two-tailed t test. n.s., not significant. Each dot represents data from one mouse. **(E)** Innervation pattern of *Cck* DMV neurons in the gastric muscularis externa and submucosal layers. Scale bar, 100 μm. **(F)** Chemogenetic activation of *Cck* DMV neurons increases gastric acid (left), pepsin (middle), and gastrin (right) secretion. Data are mean ± SEM. Unpaired two-tailed t test. **P < 0.01. Each dot represents data from one mouse. **(G)** Innervation pattern of *Grp* DMV neurons in the gastric muscularis externa and submucosal layers. Scale bar, 100 μm. **(H)** Chemogenetic activation of *Grp* DMV neurons increase gastric acid (left) secretion, with no significant change in pepsin (middle) or gastrin (right). Data are mean ± SEM. Unpaired two-tailed t test. n.s., not significant; **P < 0.01. Each dot represents one mouse. **(I–J)** Representative immunofluorescence images showing the expression of TetTox in *Cck* (I) or *Grp* (J) neurons. Scale bar, 100 μm. **(K)** Silencing *Cck* DMV neurons reduce liquid food–evoked gastric acid (left) and pepsin (right) secretion. Data are mean ± SEM. Unpaired two-tailed t test. **P < 0.01. Each dot represents one mouse. **(L)** Silencing *Grp* DMV neurons selectively reduce liquid food–evoked gastric acid (left) secretion without significantly affecting pepsin secretion (right). Data are mean ± SEM. Unpaired two-tailed t test. n.s., not significant; *P < 0.05. Each dot represents one mouse.

We expressed hM3Dq in Cre-labeled DMV neurons via AAV microinjection and activated these neurons with CNO (**Figure 5B, Figure S6A-D**). Activation of *Pdyn-Cre*^+^ DMV neurons, which preferentially innervate the gastric corpus and antrum and form terminals closely associated with Nos1^+^ myenteric neurons^23^ (**Figure 5C**), did not significantly evoke gastric secretion (**Figure 5D**). In contrast, activation of *Cck-Cre*^+^ DMV neurons robustly evoked gastric secretion (**Figure 5E–F**). Notably, activation of *Grp-Cre*^+^ DMV neurons—which preferentially innervate the gastric antrum (**Figure 5G**)—strongly increased gastric acid output but produced little to no effect on pepsin secretion (**Figure 5H**), suggesting that distinct DMV populations can differentially engage components of the secretory program.

To test whether these DMV subsets are required for physiologically evoked secretion, we silenced each population during liquid food-induced secretion. Silencing *Cck-Cre*^+^ DMV neurons reduced both gastric acid and enzyme secretion, whereas silencing *Grp-Cre*^+^ DMV neurons selectively reduced acid secretion with minimal effect on enzyme output (**Figure 5I–L**). We did not assess the role *TrpV1-Cre*^+^ DMV neurons in gastric secretion because this population did not innervate the stomach (**Figure S6E**).

### Designated enteric neurons mediate gastric secretion

Upon activation of vagal motor neurons, there are two possible mechanisms by which gastric secretion could be induced. One mechanism is a direct action of vagal motor neurons on secretory epithelial cells participating in gastric secretion (**Figure 6A, Pathway 1**). Another mechanism involves synaptic relay through local enteric neurons (**Figure 6A, Pathway 2**). To investigate the possibilities of these two mechanisms, we first characterized the terminal distribution of vagal motor neurons. We expressed tdTomato in DMV vagal motor neurons (by injecting AAV-FLEX-tdTomato into the DMV of *Chat-Cre* mice) for anterograde tracing and performed histological analysis of the stomach and small intestine. We separated the muscularis externa from the mucosa/submucosa and analyzed DMV terminals within these layers by whole-mount staining. Vagal motor neurons formed extensive terminals in the muscular layer of the stomach, with most terminals associated with myenteric ganglia^23,46^ (**Figure 6B, upper panels**). Within the submucosa, we did not observe tdTomato^+^ terminals (**Figure 6B, lower panels**). This projection pattern was also observed in the small intestine (**Figure S7A)**. In sharp contrast, *Chat-Cre*^+^ enteric neurons densely innervate the submucosa (**Figure 6C, lower panels**). These anatomical analyses argue against direct action of DMV motor neurons on secretory epithelial cells lining the mucosa but favor the second possibility: vagal motor neurons activate myenteric neurons, which in turn drive secretory epithelial cells.

**Figure 6.**
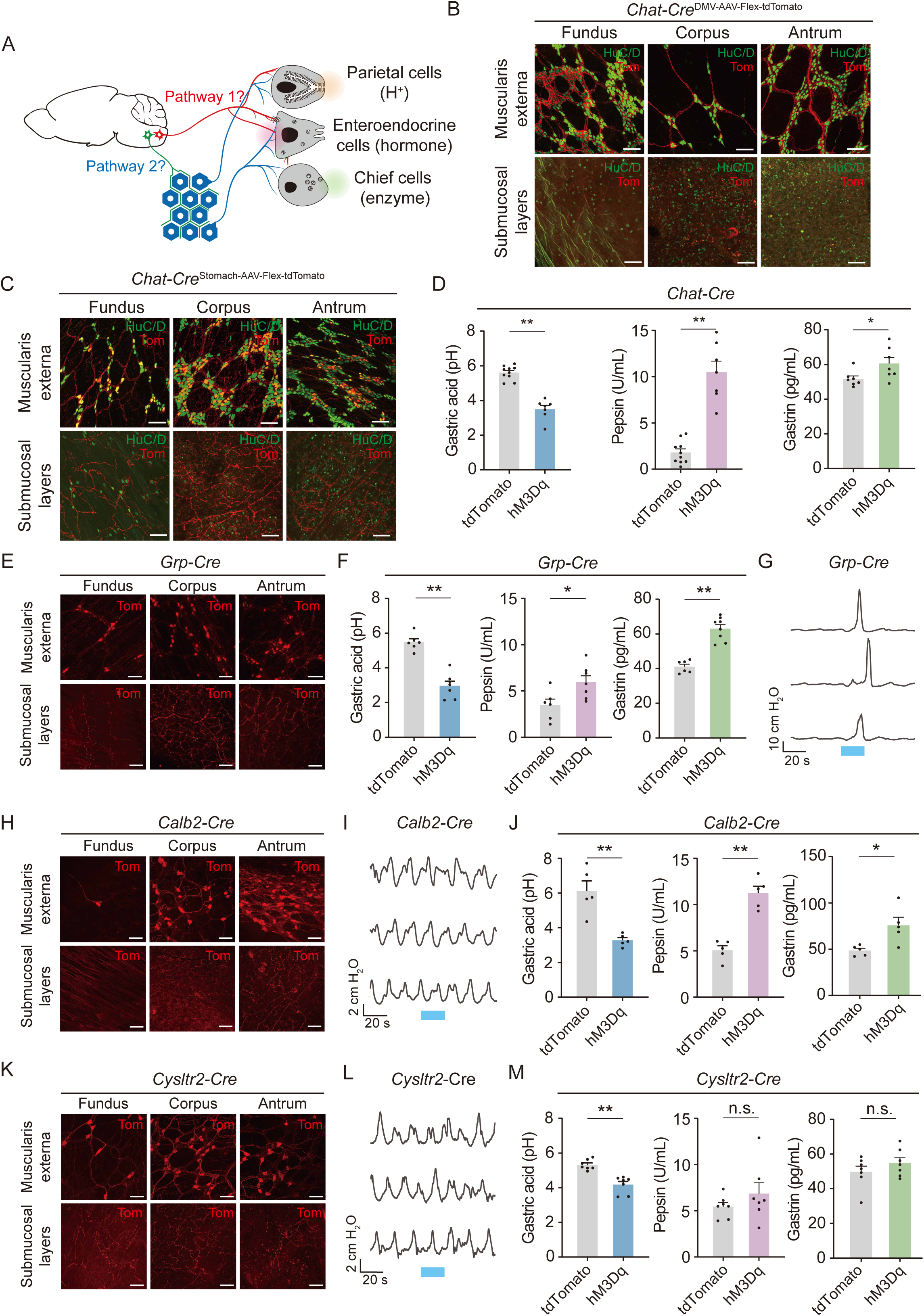
Designated enteric neurons for gastric secretion. **(A)** Schematic illustrating two vagal motor neuron–dependent pathways that regulate gastric secretion. **(B)** *Chat* DMV vagal motor neurons innervate the gastric muscularis externa but show little to no innervation of submucosal layers. HuC/D labels enteric neurons. Scale bar, 100 μm. **(C)** Distribution of *Chat* enteric neurons within the gastric muscularis externa and submucosal layers. HuC/D labels enteric neurons. Scale bar, 100 μm. **(D)** Activation of gastric *Chat* enteric neurons increase gastric acid (left), pepsin (middle), and gastrin (right) secretion. Data are mean ± SEM. Unpaired two-tailed t test. *P < 0.05; **P < 0.01. Each dot represents data from one mouse. **(E)** Distribution of *Grp* gastric neurons in the gastric muscularis externa and submucosal layers. HuC/D was used to label gastric neurons. Scale bar, 100 μm. **(F-G)** Activation of gastric *Grp* enteric neurons increase gastric secretion—acid (F, left), pepsin (F, middle), and gastrin (F, right)—and also elicits gastric motility (G). Blue light indicates the light stimulation period. Data are mean ± SEM. Unpaired two-tailed t test. *P < 0.05; **P < 0.01. Each dot represents data from one mouse. **(H)** Distribution of *Calb2* gastric neurons in the gastric muscularis externa and submucosal layers. HuC/D was used to label gastric neurons. Scale bar, 100 μm. **(I-J)** Activation of gastric *Calb2* enteric neurons increase gastric secretion—acid (H, left), pepsin (H, middle), and gastrin (H, right)—but does not significantly change gastric motility (I). Blue light indicates the light stimulation period. Data are mean ± SEM. Unpaired two-tailed t test. **P < 0.01. Each dot represents data from one mouse. **(K)** Distribution of *Cysltr2* gastric neurons in the gastric muscularis externa and submucosal layers. HuC/D was used to label gastric neurons. Scale bar, 100 μm. **(L-M)** Activation of gastric *Cysltr2* enteric neurons selectively increase gastric acid secretion (I, left) without significantly affecting pepsin (I, middle) or gastrin (I, right), and does not significantly change gastric motility (J). Blue light indicates the light stimulation period. Data are mean ± SEM. Unpaired two-tailed t test. n.s., not significant; **P < 0.01. Each dot represents data from one mouse.

If enteric neurons are indeed required for vagal motor neurons to regulate gastric secretion, silencing their activity would dampen gastric secretion evoked by either liquid food or chemogenetically induced vago–vagal reflex. To test this, we injected AAV-hSyn-Cre into the stomach of *Chat^flox/flox^*animal to block acetylcholine synthesis in enteric neurons. Littermate animals receiving control AAV virus (which do not express Cre) injections in the stomach served as a control. Compared to control animals, knocking down *Chat* expression in enteric neurons resulted in a reduction of liquid food induced gastric secretion (**Figure S7B**). In a separate cohort of *Chat^flox/flox^*mice, we inject both AAV-hSyn-Cre in the stomach and co-injected Cre and Cre-dependent hM3Dq AAVs in the nodose ganglion (for chemogenetic activation). Control animals are Littermates receiving the same AAV injections in the nodose ganglia but receiving AAV-tdTomato in the stomach. Gastric secretion induced by CNO injection (to activate stomach-innervating vagal sensory neurons) was also significantly reduced in *Chat* knocking down animals (**Figure S7C**).

Enteric neurons comprise transcriptional distinct subsets and can be subdivided into two largely non-overlapping cohorts based on neurotransmitter identity: cholinergic neurons (marked by *Chat-Cre*) and non-cholinergic subpopulations (marked by *Nos1-Cre* or *Vip-Cre*). To test their role in gastric secretion, we expressed hM3Dq with local AAV injection and chemogenetically activated cholinergic and non-cholinergic myenteric neurons. Local AAV injection led to specific gene expression in gastric enteric neurons without infecting vagal sensory neurons (**Figure S8**). Activating cholinergic neurons led to increased secretion of gastric acid, pepsin, and gastrin (**Figure 6D**), as well as stomach motility (**Figure S7E**). Conversely, activating non-cholinergic myenteric neurons with either *Nos1-Cre* or *Vip-Cre* did not induce gastric secretion (**Figure S7D**). These findings suggest that vagal motor neurons recruit cholinergic myenteric neurons, which subsequently activate the secretory epithelium and stomach motility program.

A distinct subtype of cholinergic myenteric neuron expressed gastrin-releasing peptide (Grp)^46,47^. This population stood out as a candidate enteric subtype capable of coordinating gastric functions. To test whether GRP-expressing enteric neurons are present in the stomach, we injected AAV-FLEX-tdTomato into the gastric wall of *Grp-Cre* mice and performed histological analysis. We observed robust tdTomato labeling of GRP-lineage myenteric neurons in the stomach, confirming that Grp-Cre captures a gastric enteric neuron population. These *Grp-Cre*^+^ enteric neurons extended processes toward smooth muscle and mucosal layers, consistent with the possibility that they could influence both motor and secretory outputs (**Figure 6E**). We next asked how *Grp-Cre*^+^ enteric neurons contribute to gastric physiology using chemogenetic and/or optogenetic manipulation (see Methods). Activation of *Grp-Cre*^+^ enteric neurons modulated both gastric motility and gastric secretion, indicating that this subtype can engage the motor and secretory arms of gastric control (**Figure 6F-G and Figure S7F**).

Enteric circuits are known to regulate both gastric motility and secretion, but whether these two outputs are coordinated by the same versus distinct enteric neuron subtypes remains unclear. We therefore tested two additional gastric enteric neuron populations labeled by *Cysltr2-Cre* and *Calb2-Cre*. To assess motor function, we optogenetically activated *Cysltr2* or *Calb2* enteric neurons and quantified contractile activity using an *ex vivo* gastric motility assay^48^ (Methods). In this assay, activation of either *Cysltr2* or *Calb2* neurons produced little to no change in motility compared with controls (**Figure 6I, 6L and Figure S7G-H**), suggesting that neither subtype is a major driver of the motor program under these conditions.

We then examined gastric secretory output. Chemogenetic activation of *Calb2* enteric neurons elicited a strong secretory response, including increased gastric acid output and pepsin secretion, increased gastrin levels (**Figure 6J**). On the other hand, activation of *Cysltr2* enteric neurons produced minimal (for gastric acid) or no detectable changes (pepsin and gastrin) on secretory measures (**Figure 6M**), although both *Calb2* and *Cysltr2* enteric neurons innervate the submucosa layer (**Figure 6H and K**). Together, these data identify *Calb2* enteric neurons as a potent pro-secretory module with little impact on motility, whereas *Cysltr2* neurons show limited influence on either secretion or motility in our assays. Importantly, the differential phenotypes across GRP-, Cysltr2-, and Calb2-labeled enteric neuron populations support a model in which gastric motility regulation and secretory modulation are functionally segregated across enteric neuron subtypes, rather than obligatorily co-regulated by a single enteric program.

## DISCUSSION

The vago–vagal reflex is a core parasympathetic circuit that transforms meal-derived signals into coordinated gastric responses, including secretion and motility^24,28^. Here, we provide a cell-type–resolved mechanism for how these reflex drives gastric secretion. By combining quantitative measurement of acid, pepsin, and gastrin secretion with targeted circuit perturbations, we show that stomach-innervating vagal sensory neurons are necessary and sufficient to evoke robust secretory output. We further resolve the afferent limb into functionally distinct pathways: *Glp1r* mechanosensory neurons and *Sst* antrum-innervating neurons strongly promote secretion, whereas *Oxtr* and *Vip* afferents contribute little. Mechanistically, Piezo2 in vagal afferents is required for distension-evoked secretion, linking gastric stretch to secretory reflex activation.

A longstanding question is how broadly distributed vagal sensory information is routed into specific downstream programs^22,24,25,49,50^. Gut-brain signaling shapes not only feeding behavior but also gastric emptying, motility, secretion, hormone release, and glucose homeostasis ^13,15,24,25,51,52^. Prior work has established that defined vagal afferent subtypes encode distension and nutrient signals to regulate satiety and meal termination^13,15^. Our results argue that not all these signals are obligatorily coupled to secretory output. Firstly, although activating *Oxtr*^+^ vagal sensory neurons detecting gut stretch powerfully suppress feeding (**Figure S3H**), it had little effect on gastric secretion (**Figure 3E**). Secondly, activation of second-order NTS neurons and downstream PBN pathways implicated in stomach stretch robustly altered feeding, yet it did not evoke measurable gastric secretion (**Fig. S9**), indicating functional segregation downstream of vagal mechanosensory inputs—consistent with parallel processing streams that separately engage behavioral versus digestive effectors.

Our findings also extend vagal chemosensory function beyond metabolic and feeding regulation^13^ by identifying a chemosensory pathway that directly drives secretion. *Sst* vagal sensory neurons, which preferentially innervate the stomach antral mucosa/submucosa, are involved in nutrient-evoked secretory responses, which showed stimulus selectivity, with robust secretion to carbohydrates and amino acids but comparatively weak responses to lipid stimuli. This organization suggests that vagal chemosensation can be partitioned by nutrient class to tune gastric secretory programs.

On the efferent limb, our data reveal modular parasympathetic output at two levels—within the DMV and within local enteric circuits. Recent molecular and anatomical work has defined DMV subtypes with distinct projection patterns^23^; our functional perturbations show these are functionally segregated pathways. Activating *Cck* DMV neurons broadly increased acid, pepsin, and gastrin, whereas *Grp* DMV neurons preferentially drove acid and *Pdyn* DMV neurons had little effect. Strikingly, Vagal control of secretion required an enteric relay: conditional deletion of *Chat* in gastric enteric neurons abolished vagal-evoked secretion, demonstrating that vagal motor commands are executed indirectly through cholinergic enteric intermediates. Enteric subtypes further partition outputs: *Grp* enteric neurons couple secretion to motility, *Calb2* neurons drive secretion with minimal motility effects, and *Cysltr2* neurons selectively enhance acid. These results support a labeled-line like logic for the parasympathetic pathway, analogous to the sympathetic ganglia^53^, implemented by sequential DMV-to-enteric modules that route commands to distinct gastric effectors.

## RESOURCE AVAILABILITY

### Lead contact

Requests for further information and resources should be directed to and will be fulfilled by the lead contact, Jinfei D Ni

### Data and code availability

All data required to support the conclusions of this study are provided in the paper and supplementary materials. Reagents are available from the corresponding author upon reasonable request.

## ACKNOWLEDGMENTS

We thank Songjie Lv, Yi Lv, and Yuli Jiang for advice and technical support regarding instrumental and animal maintenance, Drs. Wei Lu, Yiquan Tang and Shaoli Wang for transgenic animals. Drs. Baobin Li, Zhiyong Xie, Rongfeng Hu and Tong Ma for technical suggestions. This work is supported by grants from STI2030-Major Projects (2021ZD0203303 to J.F.N); Fund of Fudan University and Cao’ejiang Basic Research (24FCB03 to J.F.N).

## AUTHOR CONTRIBUTIONS

Conceptualization: JFN, WD, SD, KL, ZW, QT

Methodology: ZW, QT, KL, JFN, XHS, JHM, YW, LJ, BL, BX

Investigation: ZW, QT, KL, JHM, CYY

Visualization: ZW, QT,

Funding acquisition: JFN

Project administration: JFN

Supervision: JFN

Writing: JFN with the input from other authors

## STAR**Z⍰**METHODS

Detailed methods are provided in the online version of this paper and include the following:

- KEY RESOURCES TABLE
- EXPERIMENTAL MODEL AND STUDY PARTICIPANT DETAILS
- METHOD DETAILS

`o`Local AAV injection in Nodose ganglion Local Adeno-associated viral (AAV) Injections
`o`Local AAV injection in Stomach or Duodenum
`o`Subdiaphragamatic vagotomy surgery
`o`Stereotaxic injection
`o`Gastric balloon Inflation
`o`Measurement of gastric acid and pepsin
`o`Measurement of gastrin
`o`Behavioral assays
`o`Histology
`o`Ex vivo optogenetic control of gut motility
`o`Single cell RNAseq data analysis
`o`Quantification of nodose ganglion Ending Projections to the GI Tract
`o`Frozen Section Histology
`o`Behavioral and physiological assays
- QUANTIFICATION AND STATISTICAL ANALYSIS

## STAR**Z⍰**METHODS

### KEY RESOURCES TABLE

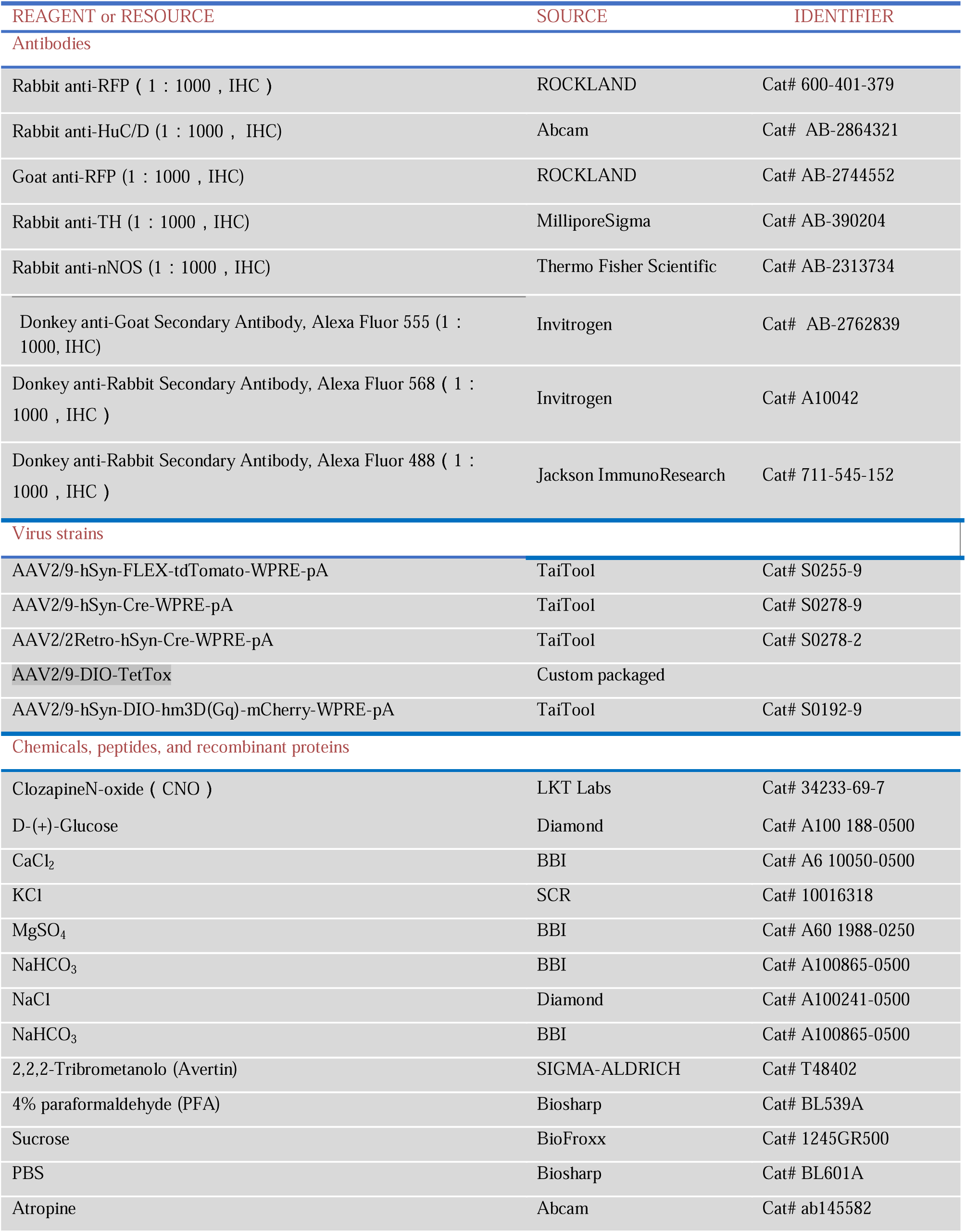
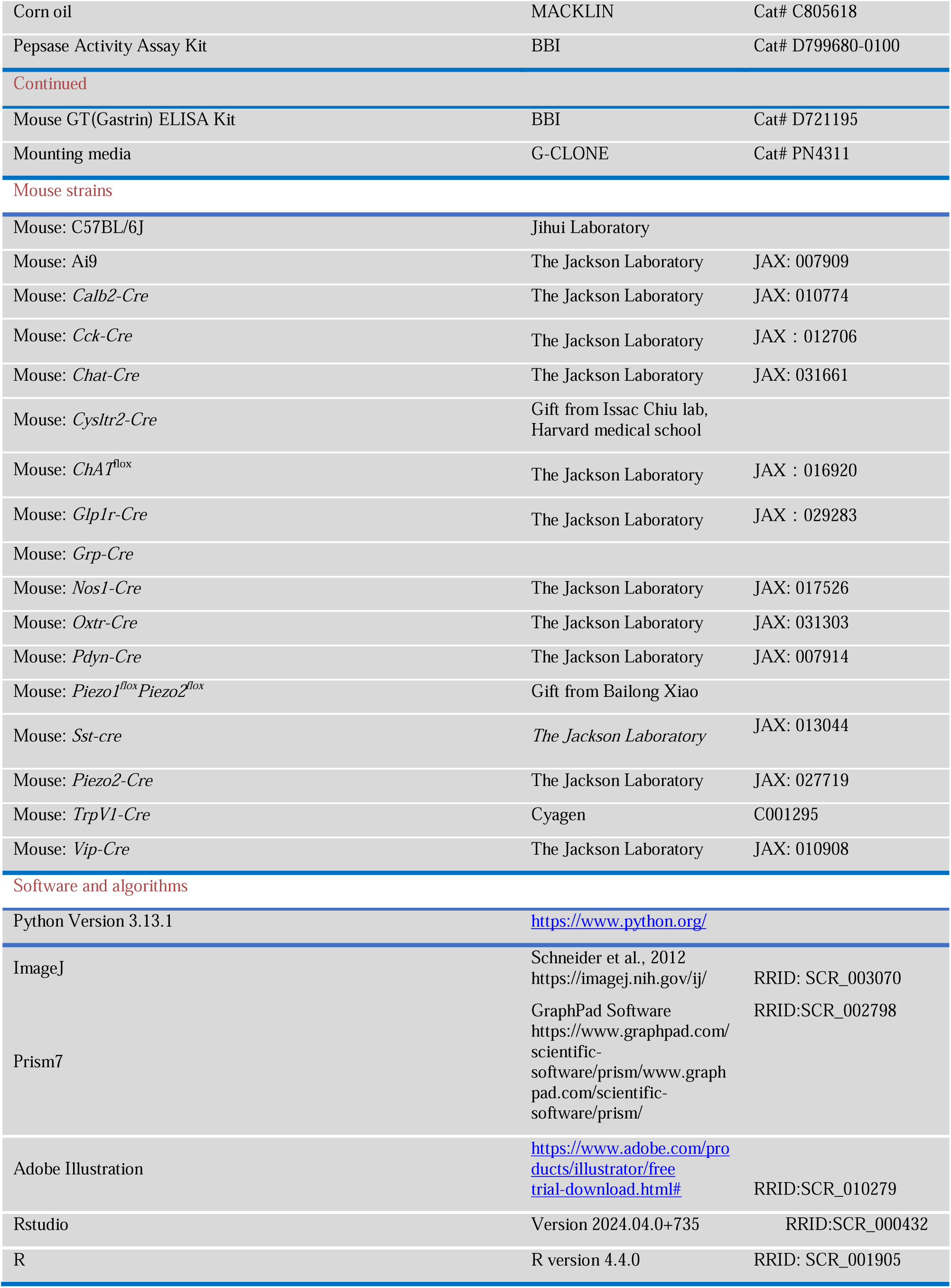

### EXPERIMENTAL MODEL AND STUDY PARTICIPANT DETAILS

Animal experiments were performed with protocols approved by the Laboratory Animal Resources Center and followed the guidelines of the Animal Experiment and Use Committee at the Shanghai Medical School of Fudan University. Mice were kept with a 12-h/12-h light/dark cycle and had *ad libitum* access to standard laboratory mouse food and water, and in a temperature-and humidity-controlled room. Both male and female mice aged 6–12 weeks were used for all experiments. The transgenic mouse lines used in this study included C57BL/6J, *Glp1r-Cre*, *Grp-Cre*, *Sst-Cre*, *Chat^flox/flox^*, *Pdyn-Cre*, *Trpv1-Cre*, *Chat-Cre*, *Nos1-Cre*, *Vip-Cre*, *Calb2-Cre*, and *Cysltr2-Cre* mice. The wild type mice (C57BL/6J) were purchased from Shanghai Jihui Laboratory Animal Care Co., Ltd. Information regarding transgenic animal strains is listed in the supplementary table.

## METHOD DETAILS

### Local AAV injection in Nodose ganglion

#### Surgical Exposure of the Vagal Ganglion

Mice were anesthetized by intraperitoneal injection of Avertin and placed supine on a surgical platform. Limbs and head were secured with adhesive tape, with the head positioned as low as possible to optimize exposure and stabilization of the cervical region. The ventral neck was dehaired, and the skin was disinfected with 75% ethanol.

A midline cervical incision (approximately 1–3 cm) was made along the sagittal plane from the mandible to the upper sternum. The bilateral submandibular glands were gently separated using fine forceps and retracted laterally to expose the trachea and carotid sheath. Under a surgical microscope, the main trunk of the vagus nerve was identified and carefully blunt-dissected away from the carotid artery. The carotid artery was then gently retracted medially toward the trachea to improve visualization of the vagus nerve trunk. Surrounding connective tissue was slowly cleared by blunt dissection to fully expose the nerve. Excessive manipulation of the vagus nerve trunk (e.g., pinching, pulling, or stretching) was avoided throughout the procedure. To further expose the nodose ganglion, the sternocleidomastoid muscle was retracted laterally using a traction hook. The hypoglossal nerve and adjacent carotid structures beneath the muscle were carefully separated by blunt dissection. The nodose (vagal) ganglion was identified as a distinct fusiform swelling along the vagus nerve trunk at this level.

#### AAV Injection

Following surgical exposure of the nodose ganglion, the ganglion was positioned for stereomicroscope-guided microinjection using a Nanoinjector (Drummond, Nanoject III) and glass micropipette. Immediately before injection, AAV was mixed with 5× Fast Green (4:1, v/v) to visualize delivery and was kept on ice until use. The AAV/Fast Green mixture was then loaded into the pipette tip. Under direct visualization, the pipette tip was advanced into the nodose ganglion and 200 nL of virus was delivered at 2 nL/s. After completion of injection, the pipette was left in place briefly and then withdrawn slowly to minimize reflux. The incision was closed with sutures and the wound was disinfected with iodophor.

### Local AAV injection in Stomach or Duodenum

Injection produce was performed as previously described. Under a stereomicroscope, the stomach or duodenum was exposed through a midline abdominal incision. A total volume of 5 µL AAV was injected into the gastric or duodenal wall using a nanoinjector at a rate of 50 nL/s, with 50 nL per injection site. Injection sites were evenly distributed to cover the entire tissue surface. During injection, any viral leakage was immediately removed using sterile cotton swabs to prevent unintended viral spread. All behavioral and histological experiments were performed at least 4 weeks after AAV injection.

### Subdiaphragamatic vagotomy surgery

Mice were anesthetized and placed supine on a warmed surgical surface. After abdominal disinfection, a ∼2 cm midline laparotomy was made to expose the stomach and distal esophagus. Under a dissecting microscope, the anterior and posterior subdiaphragmatic vagal trunks/branches running along the esophagus and projecting to the stomach were identified, carefully separated from surrounding connective tissue, and transected bilaterally. A ∼2 mm segment was removed from each branch to create a gap and minimize reinnervation. The abdominal muscle wall and skin were closed in layers with sutures. Mice were maintained on a heating pad until fully recovered from anesthesia and then returned to their home cages. All procedures were performed under aseptic conditions.

### Stereotaxic injection

Mice were anesthetized and secured in a stereotaxic apparatus (RWD, China). For DMV, viral vectors were injected via Nanoinjector and glass micropipette at a rate of 60 nL/min, with a standard volume of 15 nL per site.

The dorsal motor nucleus of the vagus (DMV) was targeted at three distinct anteroposterior levels relative to bregma. The coordinates were as follows: rostral DMV (−7.15 mm AP; ±0.25 mm ML; −4.40 mm DV), intermediate DMV (−7.50 mm AP; ±0.25 mm ML; −4.50 mm DV), and caudal DMV (−7.90 mm AP; ±0.20 mm ML; −4.55 mm DV). Injections were performed vertically.

The nucleus tractus solitarius (NTS) was targeted at the following coordinates relative to bregma: −7.40 mm anteroposterior (AP), ±0.65 mm mediolateral (ML), and −4.40 mm dorsoventral (DV).

The parabrachial nucleus (PBN) was targeted using the following coordinates relative to bregma:-5.15 mm anteroposterior (AP); ±1.35 mm mediolateral (ML);-2.10 mm dorsoventral (DV). Injections were performed vertically.

Mice were allowed to recover for 4 weeks to ensure robust viral expression before the commencement of experiments.

### Gastric balloon Inflation

Mice were anesthetized with Avertin, a balloon was then inserted into the stomach via the esophagus and subjected to mechanical distension by filling the balloon with specific volume of water. After 30 minutes, the stomach was rapidly excised, and gastric acid and pepsin were measured following the aforementioned procedure.

### Measurement of gastric acid and pepsin

Animals are fasted for 8–12 hours and water deprivation for around 6 hours prior to the experiment.

For chemogenetic experiments, mice received an intraperitoneal injection of clozapine-N-oxide (CNO, 3 mg/kg), and gastric acid and pepsin levels were measured 1 hour after injection. Briefly, mice were rapidly anesthetized with isoflurane and sacrificed. The following procedure is used to collect stomach content: for CNO chemogenetic experiment, 100 μL of saline was perfused into the stomach via the cardia to wash out luminal acid. For A small incision was made in the nonglandular stomach with fine scissors, and the gastric contents were squeezed out into a 1.5 mL centrifuge tube, which was then centrifuged at 10,000 rpm and 4°C for 10 minutes. The supernatant was collected for pH measurement using a micro-pH meter (LabSen241-3A micro-pHmeter, China).

For nutrient-evoked stimulation experiments, mice were orally gavaged with 400 µL of nutrient solution, and gastric acid and pepsin levels were assessed 30 min after gavage. Briefly, mice were rapidly anesthetized with isoflurane and sacrificed. The stomach was acutely excised. A small incision was made in the nonglandular stomach with fine scissors, and the gastric contents were squeezed into a 1.5 mL centrifuge tube, which was then centrifuged at 10,000 rpm and 4°C for 10 minutes. The supernatant was collect for pH measurement using a micro-pH meter (LabSen241-3A micro-pHmeter, China).

For mechanical-evoked stimulation experiments. Mice were anesthetized with Avertin, a mini balloon was inserted into the stomach along the esophagus, and then the stomach was subjected to balloon distension with a volume of 300 μL for 30 minutes. Finally, the mouse’s stomach was rapidly excised, and 100 μL of saline was perfused into the stomach via the cardia wash out luminal acid. For A small incision was made in the nonglandular stomach with fine scissors, and the gastric contents were squeezed into a 1.5 mL centrifuge tube, which was then centrifuged at 10,000 rpm and 4°C for 10 minutes. The supernant was collect for pH measurement using a pH meter.

After measuring the pH, the contents in the 1.5 mL centrifuge tube were centrifuged again at 10,000 rpm and 4 °C for 10 minutes, the supernatants were collected and then assayed for pepsin using a Pepsase Activity Assay Kit (Sangon# D799680-0100), following the manufacturer’s instructions.

### Measurement of gastrin

For gastrin measurements, mice underwent 1 h of chemogenetic activation and were then sacrificed. Blood (∼ 600 µL) was collected by cardiac puncture and centrifuged at 1,000 × g, 4°C, for 20 min to isolate serum. Serum gastrin levels were quantified using a Mouse GT (Gastrin) ELISA Kit (Sangon Biotech, D721195) according to the manufacturer’s instructions.

### Behavioral assays

Mice of either sex (8-14 weeks old) were tested ≥ 4 weeks post-viral injection. A magnetically mounted, 3D-printed custom feeder insert was placed in the test cage to hold pellets, allow free access to food, and prevent spillage into the bedding. The insert could be removed at any time for weighing, enabling precise quantification of intake. Before testing, mice were single-housed in a test cage identical to the home cage for 3 days to habituate and learn to eat from the insert. Two feeding paradigms were used.

Food intake was assessed under fasted conditions. Mice were food deprived overnight prior to testing. Sessions began at 20:00 (local time). CNO or saline (vehicle) was administered 15 min before session onset, pre-weighed food was placed in the home cage and food intake was measured by weighing the remaining food at 1-h intervals for a total duration of 3 h, and food consumption was calculated accordingly.

For the ad libitum feeding behavior test, mice were not food-deprived prior to the experiment. Sessions began at 20:00 (local time). CNO or saline (vehicle) was administered 15 min before session onset, pre-weighed food was placed in the home cage. Food intake was measured at 1-h intervals for 3 h, as described above.

### Histology

#### Tissue preparation

For gastrointestinal tissues including muscularis externa and submucosa layers, the tissues of anesthetized mice were dissected and placed in cold PBS, cutting along the side of the mesentery. Luminal contents were removed by rinsing with ice-cold phosphate-buffered saline (PBS). Tissues were then pinned flat in Sylgard™-coated dishes and fixed in 4% paraformaldehyde (PFA) at room temperature for 90 min. Following fixation, the muscular and mucosal layers were carefully separated using fine forceps for whole-mount staining.

For nodose ganglion whole-mount immunofluorescence, mice were anesthetized and perfused with 20 mL PBS followed by 20 mL of ice-cold 4% PFA. Nodose ganglion were dissected and post-fixed in ice-cold PFA for 1 hour. The tissues were then processed for whole-mount immunofluorescence staining.

#### Whole-mount immunofluorescence

The fixed gut tissue and/or nodose ganglion were washed three times for 10 minutes each in PBS buffer. The samples were then blocked in a PBS buffer containing 5% normal donkey serum, 0.3% Triton X-100 (blocking solution), and gently shaking for 1 hour at room temperature. Primary antibodies were dissolved in the blocking solution at specified concentrations (Goat anti-RFP 1:1000, Rabbit anti-HuC/D 1:1000, Rabbit anti-nNOS 1:1000). After 24 hours of incubation with primary antibodies solution at 4°C, samples were washed three times in PBST buffer. The samples were then incubated with secondary antibodies, also dissolved in the blocking solution (1:1000), overnight at 4°C. The secondary antibodies included donkey anti-rabbit Alexa Fluor 488, donkey anti-Goat Alexa Fluor 555. After incubation, samples were washed three times in PBST buffer and mounted in mounting media containing DAPI (G-CLONE).

#### Tissue sectioning and Immunofluorescence

Mice were anesthetized with isoflurane and perfused with 20 mL PBS followed by 20 mL of ice-cold 4% PFA. The brain and nodose ganglion were quickly removed intact and fixed in 4% PFA at 4°C for 24 hours. The surface liquid was blotted with filter paper, and the tissues were then subjected to gradient dehydration in 15% and 30% sucrose solutions at 4°C. After complete dehydration, OCT was used for tissue embedding.

Coronal brain sections, 50 µm thick, were prepared using a Leica freezing microtome (CM1950) and collected in PBS buffer. Free-floating brain sections were washed three times in PBS buffer to remove the OCT. Sections were permeabilized using PBST three times. Sections were blocked in blocking solution at room temperature for 1 hour and then incubated with the primary antibody diluted (1:1000) in blocking solution overnight at 4°C. The primary antibody used was Rabbit anti-RFP or Goat anti-RFP. After primary antibody incubation, sections were rinsed three times in clean PBST before incubating with secondary antibodies diluted (1:1000) in blocking solution for 2 hours at room temperature. The secondary antibodies included Alexa Fluor 568 (Donkey anti-Rabbit) or Alexa Fluor 555 (Donkey anti-Goat). Sections were washed with PBST six times and then mounted using mounting media containing DAPI.

All fluorescent images, whether single plane or z-stacks, were captured using an Olympus confocal microscope (FV-3000) and processed using ImageJ analysis software (ImageJ 1.53t).

#### *Ex vivo* optogenetic control of gut motility

Ex vivo gastric motility recordings were performed as previously described^48^. Briefly, mice were anesthetized and the stomach was rapidly excised (with about about 5 mm esophagus attached to the cardiac end) and rinsed in oxygenated Krebs buffer maintained at 37°C. The tissue was transferred to a custom organ chamber perfused with continuously oxygenated (95% O /5% CO) Krebs buffer maintained at 37°C.

The residual esophagus attached to the cardiac end of the stomach was ligated with suture to create a sealed end. The pyloric (distal) end was cannulated with a silicone tube filled with Krebs buffer and connected to a pressure transducer (TSD104A, BIOPAC Systems, USA). Intragastric pressure fluctuations generated by spontaneous or evoked contractions were converted to electrical signals and recorded with an MP160 data acquisition system (BIOPAC Systems, USA).

For optogenetic experiments, an optical fiber was positioned ∼0.5 cm above the indicated segment of the ex vivo stomach preparation. Tissue was stimulated with 405-nm light delivered by an Inper Smart Light Source while intragastric pressure signals were recorded simultaneously. Light stimulation was applied with the following parameter: 20 s length, 10 Hz, 10 ms pulse width, with 90-s inter-train intervals.

### Single cell RNAseq data analysis

Single-cell RNA sequencing data of the nodose ganglion were obtained from GSE192987 (8 datasets). Data processing and downstream analyses were performed using Seurat following standard workflows. Cells expressing fewer than 500 genes and genes detected in fewer than 10 cells were removed. The 8 datasets were integrated using the IntegrateData function to correct for batch effects.

Gastric neurons were identified by selecting cells with QZ1 > 0. These neurons were clustered using Seurat (dims = 1:30, resolution = 0.3). Cells were annotated based on expression of genes of interest (expression > 0.5), and the overlap between annotated subsets was quantified. Finally, pairwise differential gene expression analysis was performed between clusters (P-value < 0.01, fold change > 2)

### Quantification of nodose ganglion Ending Projections to the GI Tract

The distribution of vagal sensory endings across the gastrointestinal (GI) tract was quantified from whole-mount immunofluorescence images using Fiji (ImageJ). The small intestine was subdivided into duodenum, jejunum, and ileum, and the large intestine into colon and rectum. For each mouse, images were acquired from three randomly selected fields in the stomach and five randomly selected fields per intestinal segment.

For each field of view, vagal fiber density was calculated as: fiber density = (area occupied by vagal fibers) / (total imaged area). The mean fiber density across all fields within a given segment was used as the representative density for that segment in each mouse. Data were collected from three mice.

## QUANTIFICATION AND STATISTICAL ANALYSIS

All statistical analyses were performed in GraphPad Prism 7. Data are presented as mean ± SEM, as indicated in the figure legends. Sample sizes were not determined by a priori power calculations; instead, they were chosen based on animal availability and prior studies in the field. Statistical tests are specified in the corresponding figure legends and include unpaired two-tailed t tests, one-way ANOVA with Tukey’s multiple-comparisons test, and two-way ANOVA with Sidak’s multiple-comparisons test. The sample size (n) refers to the number of animals per group and is indicated in each graph and/or legend.

## SUPPLEMENTAL INFORMATION

**Document S1. Figures S1–S9 (this is the main PDF)**

## SUPPLEMENTAL FIGURES

**Figure S1.**
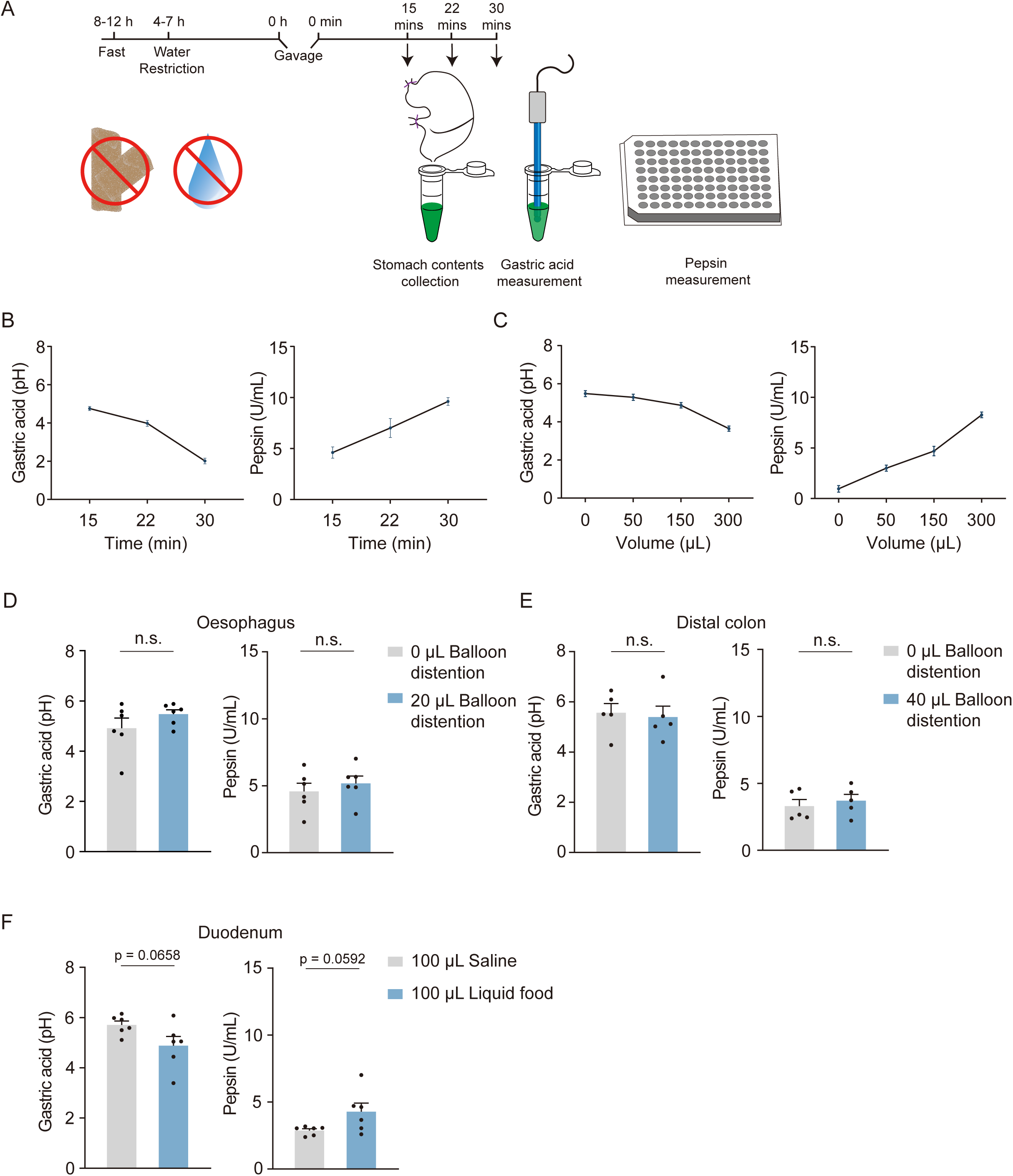
Stimulus modality-dependent regulation of gastric secretion. (A) Schematic of the experimental timeline and workflow used to quantify gastric acid and pepsin secretion. (B) Time course of gastric acid and pepsin secretion following liquid meal stimulation. Data are mean ± SEM; n = 5 mice per group. (C) Dose–response of gastric acid and pepsin secretion evoked by balloon distension. Data are mean ± SEM; n = 7 mice per group. (D) Esophageal balloon distension (20 µL) does not significantly alter gastric acid (left) or pepsin (right) secretion. Data are mean ± SEM. Unpaired two-tailed t test. n.s., not significant. Each dot represents one mouse. (E) Distal colonic balloon distension (40 µL) does not significantly alter gastric acid (left) or pepsin (right) secretion. Data are mean ± SEM. Unpaired two-tailed t test. n.s., not significant. Each dot represents one mouse. (F) Duodenal infusion of a liquid meal (100 µL) elicits a modest increase in gastric acid (left) and pepsin (right) secretion. Data are mean ± SEM. Unpaired two-tailed t test. n.s., not significant. Each dot represents one mouse.

**Figure S2.**
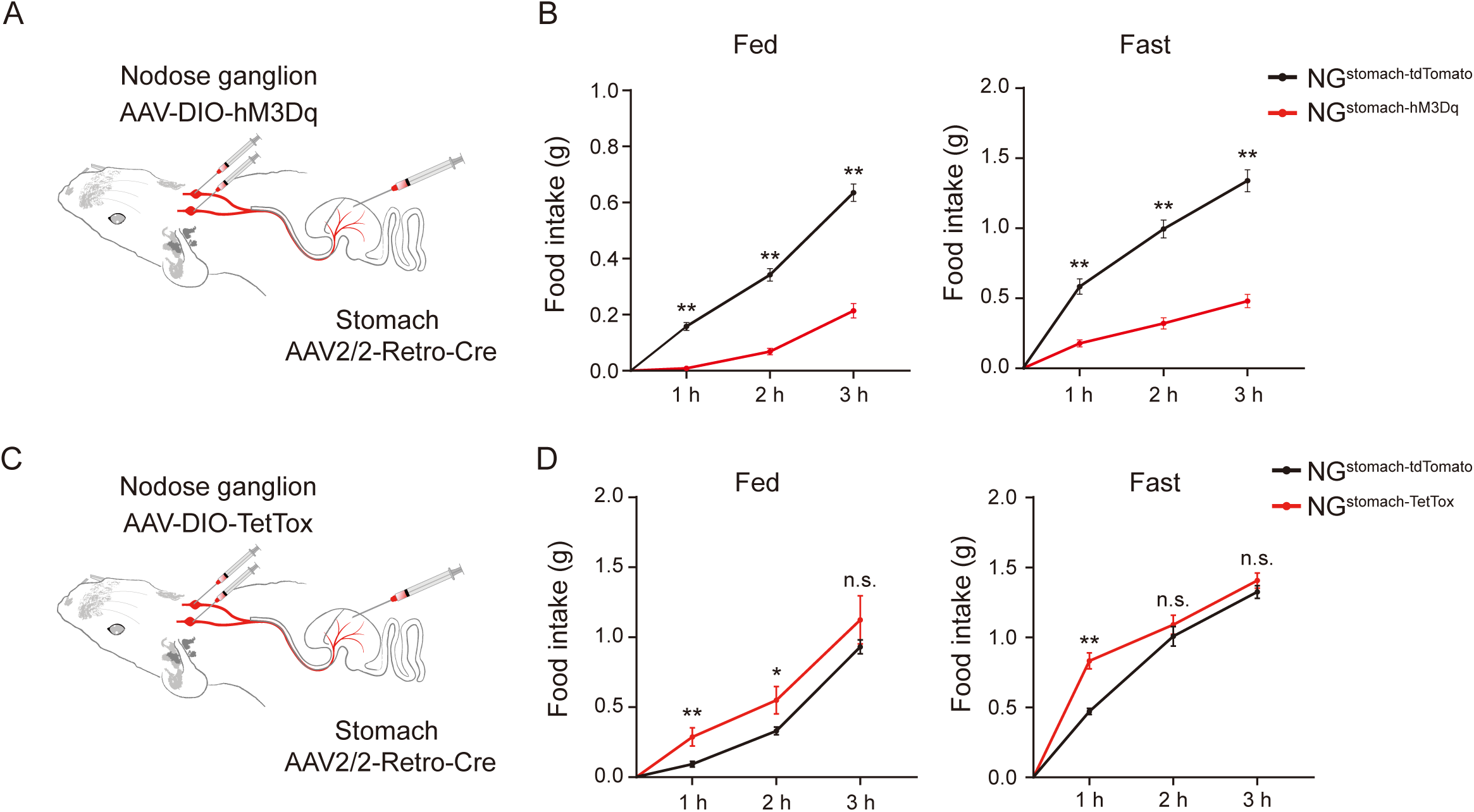
Specific manipulation of stomach-innervating vagal sensory neurons alters feeding behavior in mice. (A) Schematic of chemogenetic targeting of stomach-innervating nodose ganglion neurons (left). Chemogenetic activation of these neurons suppresses food intake in mice (right). Data are mean ± SEM. Student’s t test. **P < 0.01. Each dot represents one mouse; n = 10 mice per group. (B) Schematic of tetanus toxin–mediated synaptic silencing of stomach-innervating nodose ganglion neurons (left). Silencing these neurons increases food intake in mice (right). Data are mean ± SEM. Student’s t test. **P < 0.01. Each dot represents one mouse; n = 10 mice per group.

**Figure S3.**
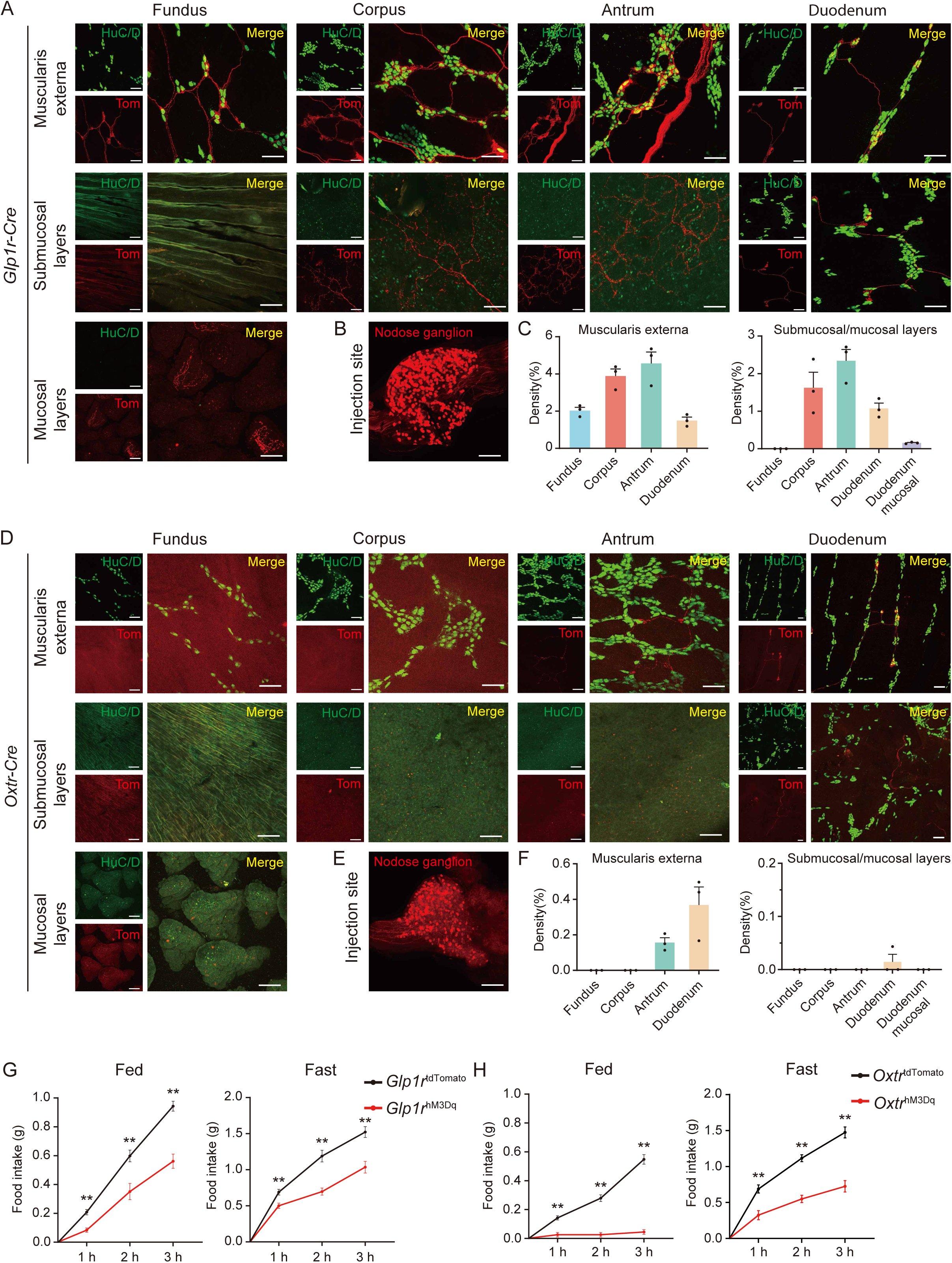
Activation of Glp1r and Oxtr vagal sensory neurons suppress food intake in mice. (A) Whole-mount immunostaining showing the distribution of Glp1r vagal sensory neuron terminals in the stomach and duodenum. HuC/D labels enteric neurons. Scale bar, 100 µm (B) Whole-mount immunostaining showing the Glp1r vagal sensory neurons infected by AAV. Scale bar, 100 µm. (C) Quantification of Glp1r vagal sensory neuron terminal density across gastrointestinal regions. Data are mean ± SEM. Each dot represents one mouse. (D) Whole-mount immunostaining showing the distribution of Oxtr vagal sensory neuron terminals in the stomach and duodenum. HuC/D labels enteric neurons. Scale bar, 100 µm. (E) Whole-mount immunostaining showing the Oxtr vagal sensory neurons infected by AAV. Scale bar, 100 µm. (F) Quantification of Oxtr vagal sensory neuron terminal density across gastrointestinal regions. Data are mean ± SEM. Each dot represents one mouse. (G) Chemogenetic activation of Glp1r vagal sensory neurons reduce food intake. Data are mean ± SEM. Student’s t test. n.s., not significant; *P < 0.05; **P < 0.01. n = 7 mice per group. (H) Chemogenetic activation of Oxtr vagal sensory neurons reduce food intake. Data are mean ± SEM. Student’s t test. n.s., not significant; **P < 0.01. n = 9 mice per group.

**Figure S4.**
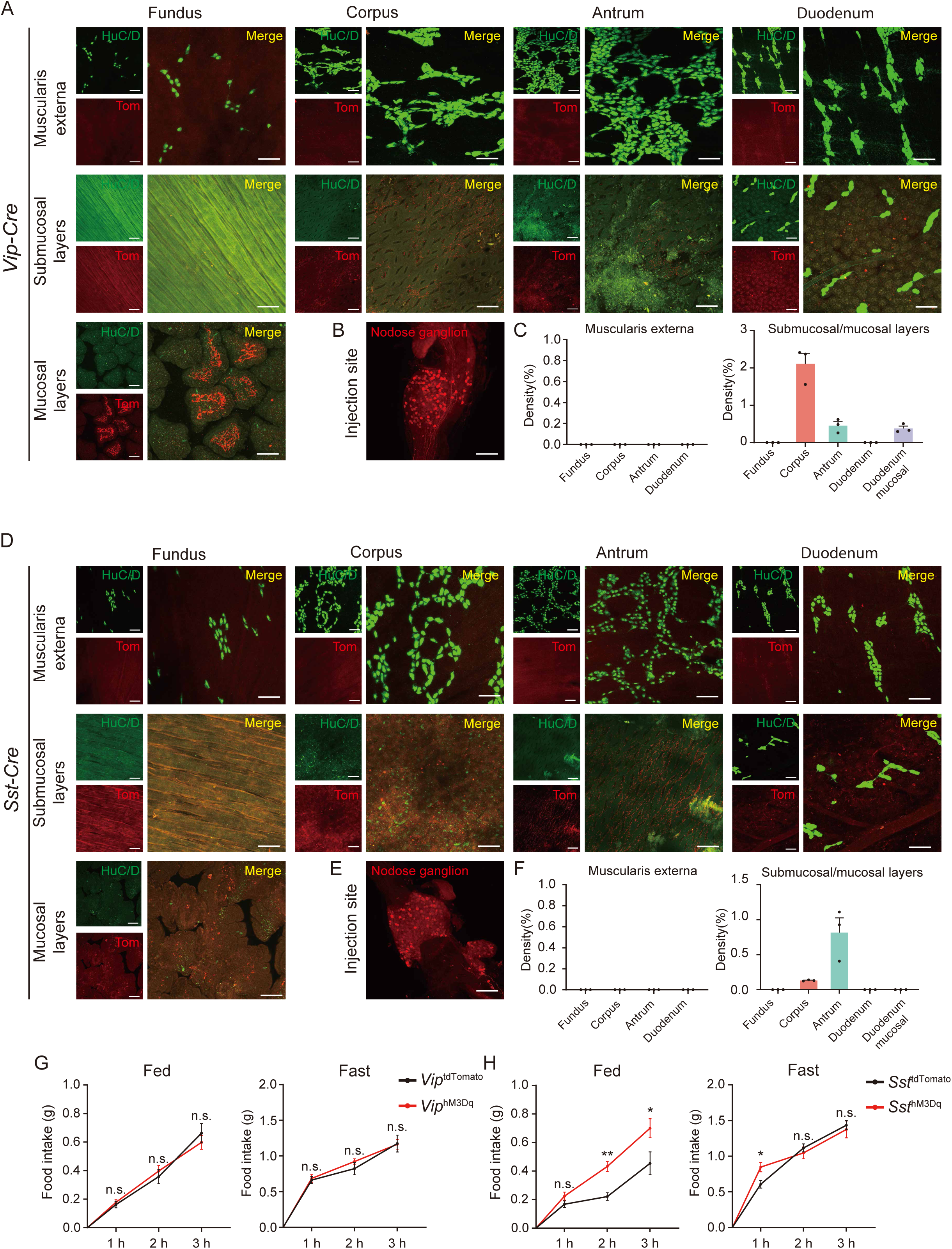
Activation of Vip**O** and Sst**O** vagal sensory neurons differentially modulate food intake in mice. (A) Whole-mount immunostaining showing the distribution of Vip vagal sensory neuron terminals in the stomach and duodenum. HuC/D labels enteric neurons. Scale bar, 100 µm. (B) Whole-mount immunostaining showing the Vip vagal sensory neurons infected by AAV. Scale bar, 100 µm. (C) Quantification of Vip vagal sensory neuron terminal density across gastrointestinal regions. Data are mean ± SEM. Each dot represents one mouse. (D) Whole-mount immunostaining showing the distribution of Sst vagal sensory neuron terminals in the stomach and duodenum. HuC/D labels enteric neurons. Scale bar, 100 µm. (E) Whole-mount immunostaining showing the Sst vagal sensory neurons infected by AAV. Scale bar, 100 µm. (F) Quantification of Sst vagal sensory neuron terminal density across gastrointestinal regions. Data are mean ± SEM. Each dot represents one mouse. (G) Chemogenetic activation of Vip vagal sensory neurons does not significantly alter food intake. Data are mean ± SEM. Student’s t test. n.s., not significant. n = 10 mice per group. (H) Chemogenetic activation of Sst vagal sensory neurons increase food intake. Data are mean ± SEM. Student’s t test. n.s., not significant; *P < 0.05. n = 8 mice per group.

**Figure S5.**
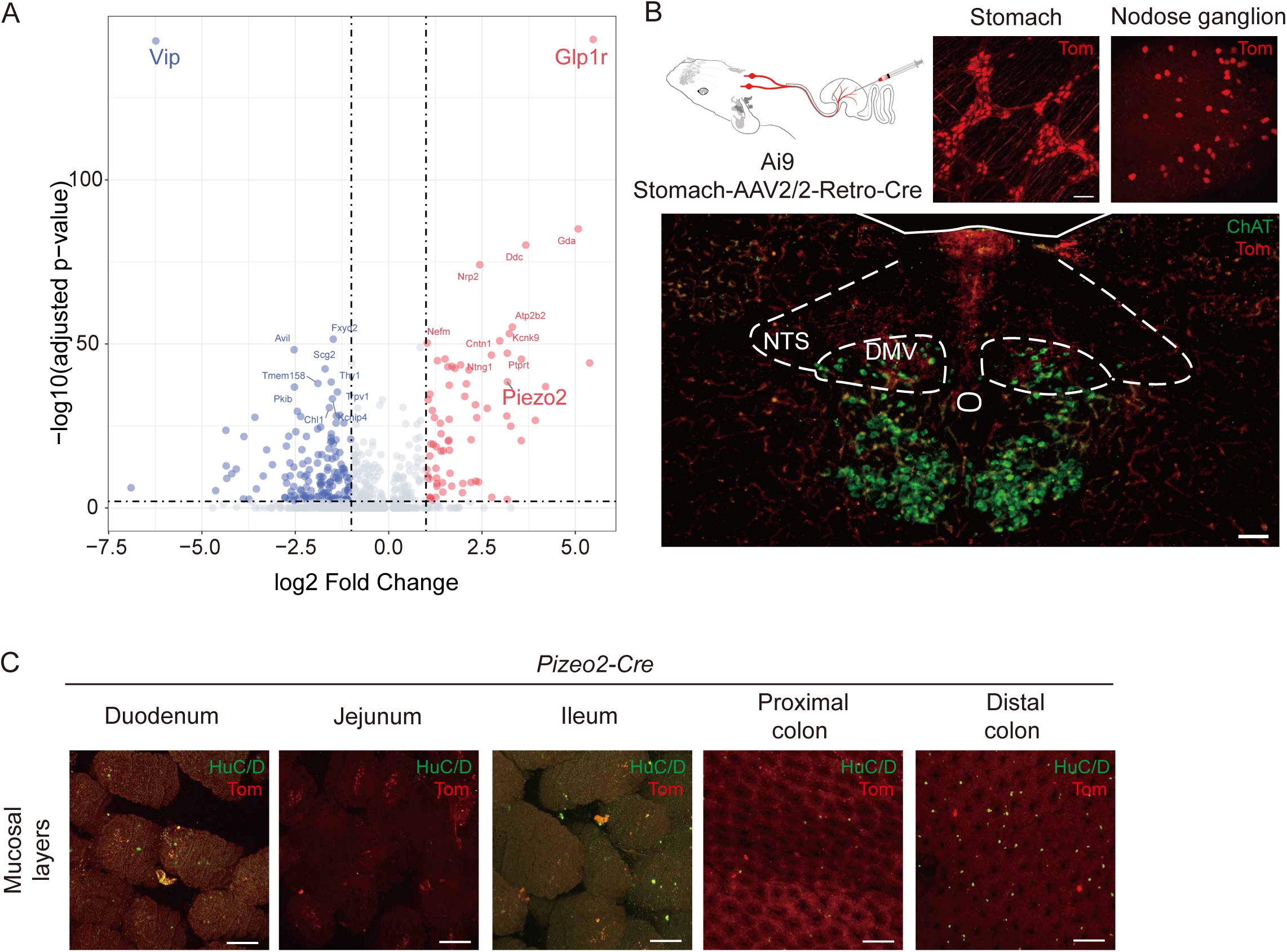
Piezo2 expression in vagal sensory neurons. (A) Differentially expressed genes between Vip and Glp1r vagal sensory neuron populations. (B) Whole-mount staining showing labeled vagal sensory neurons and local enteric neurons by retrograde AAV injection in the stomach wall. Scale bar, 100 µm. (C) Immunofluorescence showing the absence of Piezo2 vagal sensory neuron terminals in the mucosal layers of the small intestine and colon. Scale bar, 100 µm.

**Figure. S6.**
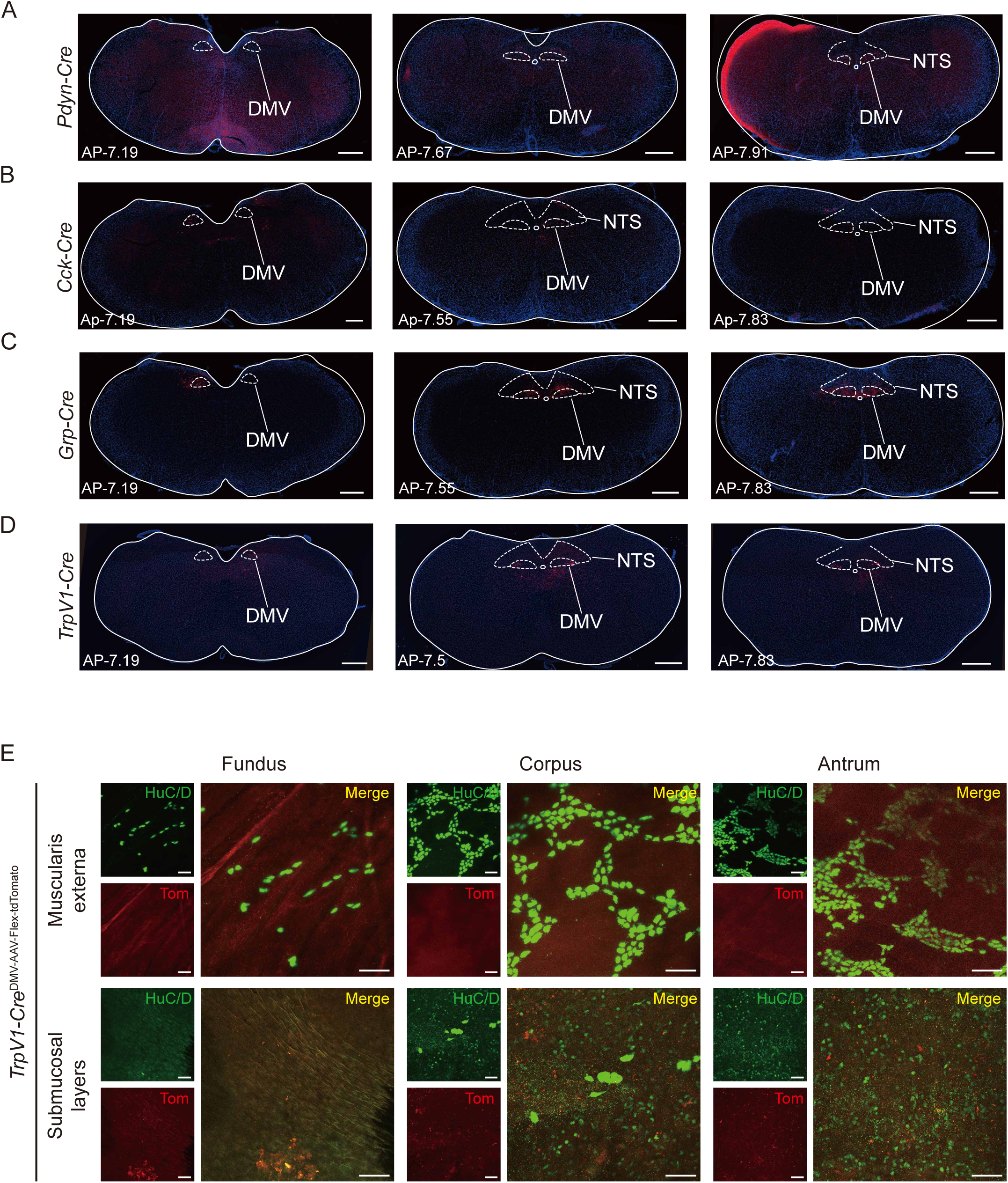
Distribution of distinct neuronal subtypes in the DMV. (**A-D**) Immunostaining of brainstem sections showing the localization of Cck (A), Pdyn (B), Grp (C), and Trpv1 (D) neurons in the dorsal motor nucleus of the vagus (DMV). Numbers on the left indicate the anterior–posterior coordinates relative to bregma. Scale bar, 500 µm (**E**) Whole-mount staining showing the absence of TrpV1^+^ vagal motor neuron terminals in the stomach. HuC/D was used to label gastric neurons. Scale bar, 100 μm.

**Figure S7.**
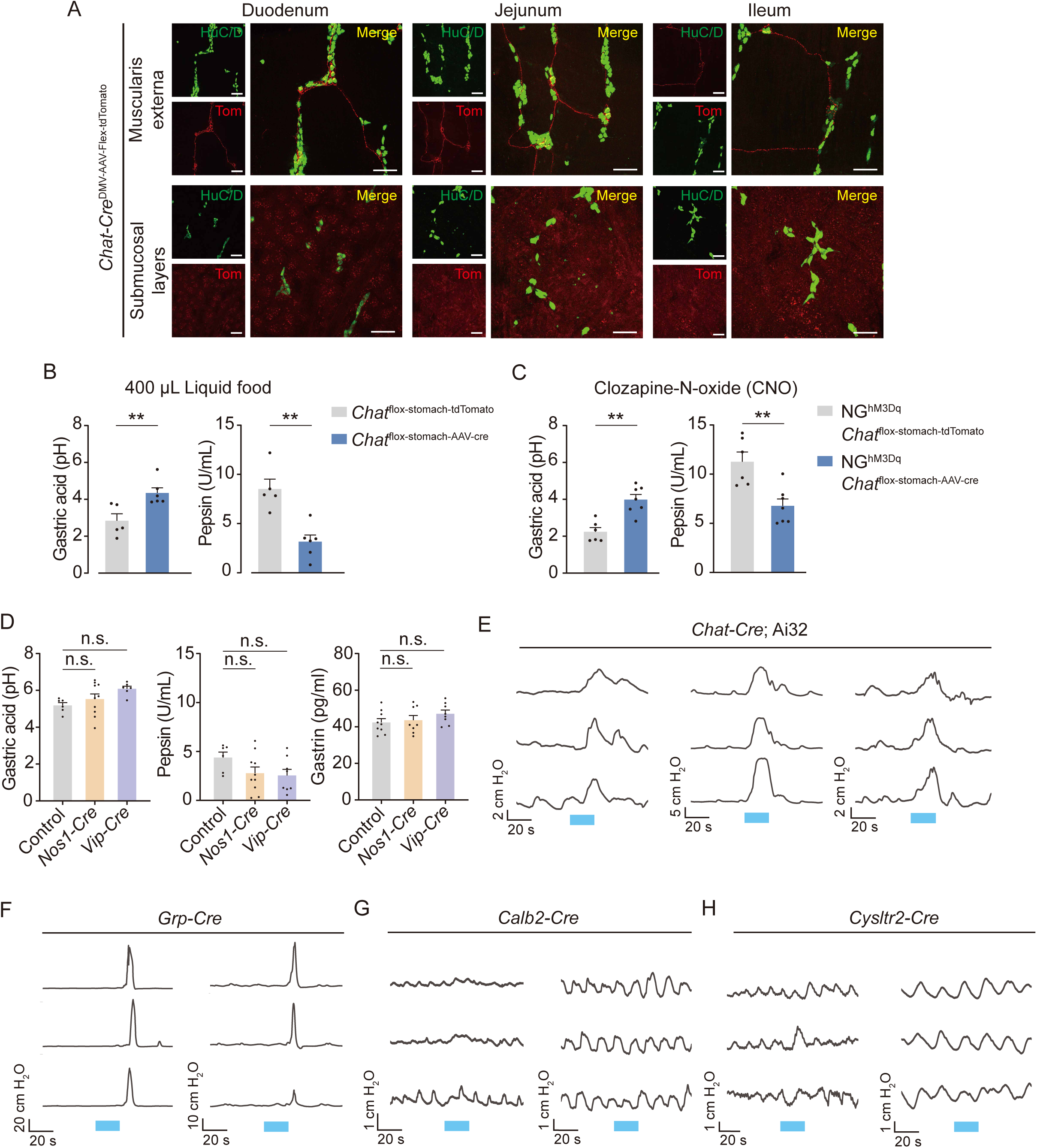
Enteric cholinergic neurons in the stomach promote gastric secretion and motility. (A) Wholemount Immunostaining showing the distribution of terminals from *Chat* neurons originating in the dorsal motor nucleus of the vagus (DMV). Scale bar, 100 µm. (B) Conditional deletion of Chat in stomach enteric neurons reduces liquid meal–evoked secretion of gastric acid (left) and pepsin (right). Data are mean ± SEM. Unpaired two-tailed t test. n.s., not significant. Each dot represents one mouse. (C) Conditional deletion of Chat in stomach enteric neurons reduces vagal sensory neuron activation–evoked secretion of gastric acid (left) and pepsin (right). Data are mean ± SEM. Unpaired two-tailed t test. n.s., not significant. Each dot represents one mouse. (D) Activation of non-cholinergic enteric neurons does not significantly alter secretion of gastric acid (left), pepsin (middle), or gastrin (right). Data are mean ± SEM. One-way ANOVA with Tukey’s multiple-comparisons test. n.s., not significant. Each dot represents one mouse. (**E-H**) Modulation of gastric motility by optogenetic activation of indicated enteric neurons.

**Fig. S8.**
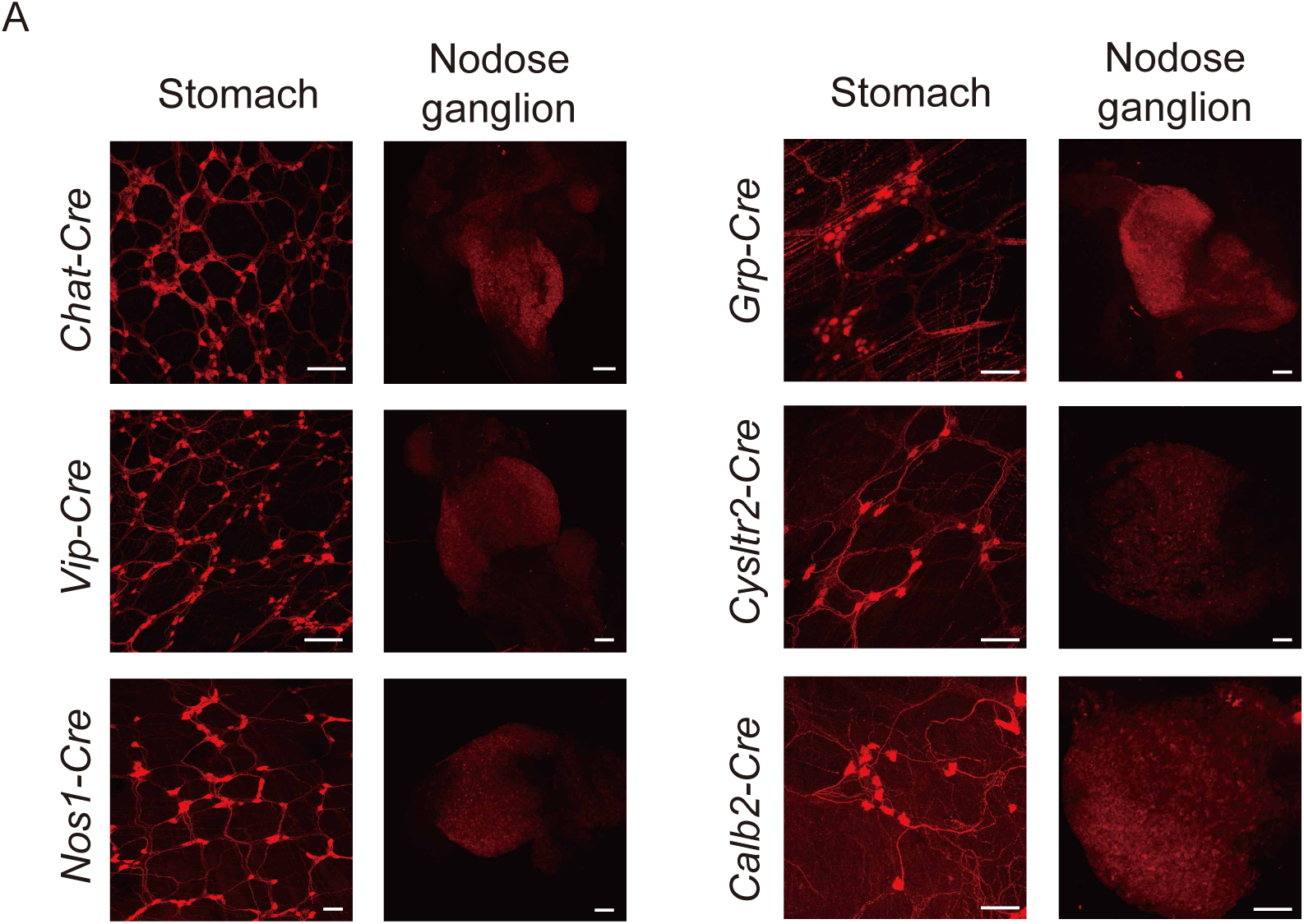
Verification of local AAV transfection in the stomach **(A)** Whole-mount immunostaining of stomach and nodose ganglion showing the infection of AAV-dio-hM3Dq in the stomach but not in the vagal sensory neurons, Scale bar, 100 µm.

**Figure S9.**
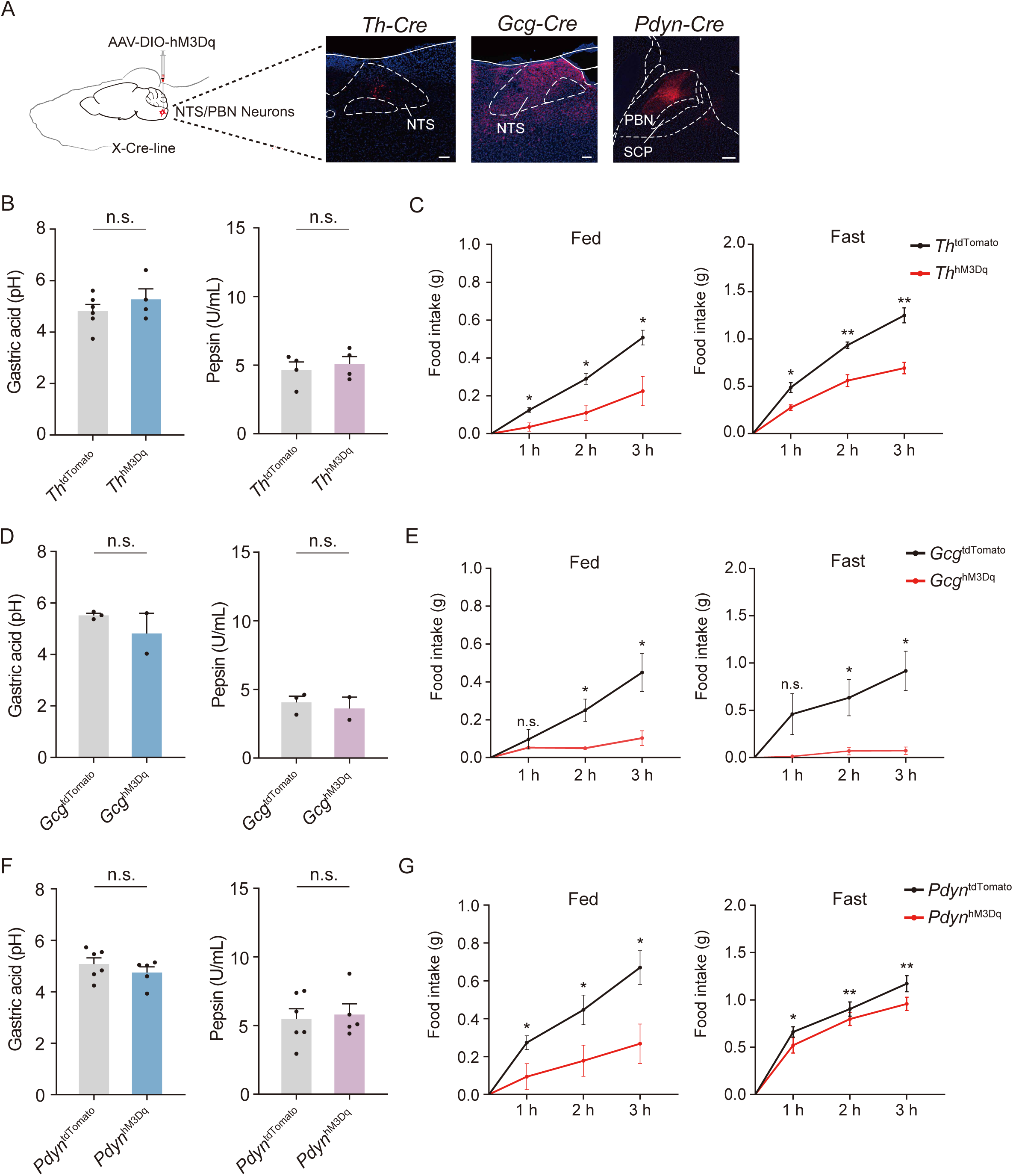
Manipulation of neuronal subtypes in the NTS and PBN differentially affects gastric secretion and feeding behavior. (B) Schematic of chemogenetic targeting in the nucleus of the solitary tract (NTS; Th or Gcg neurons) and the parabrachial nucleus (PBN; Pdyn neurons). Scale bar, 100 µm. (C) Chemogenetic activation of Th NTS neurons does not significantly alter gastric acid (left) or pepsin (right) secretion. Data are mean ± SEM. Unpaired two-tailed t test. n.s., not significant. Each dot represents one mouse. (D) Chemogenetic activation of Th NTS neurons reduces food intake. Data are mean ± SEM. Student’s t test. *P < 0.05; **P < 0.01. n = 4 mice per group. (E) Chemogenetic activation of Gcg NTS neurons does not significantly alter gastric acid (left) or pepsin (right) secretion. Data are mean ± SEM. Unpaired two-tailed t test. n.s., not significant. Each dot represents one mouse. (F) Chemogenetic activation of Gcg NTS neurons does not significantly alter food intake. Data are mean ± SEM. Student’s t test. n.s., not significant. n = 3 mice per group. (G) Chemogenetic activation of Pdyn PBN neurons does not significantly alter gastric acid (left) or pepsin (right) secretion. Data are mean ± SEM. Unpaired two-tailed t test. n.s., not significant. Each dot represents one mouse. (H) Chemogenetic activation of Pdyn PBN neurons reduces food intake. Data are mean ± SEM. Student’s t test. n.s., not significant. n = 5 mice per group.

